# Two functionally distinct amyloid-β intermediates link aggregation pathways to pathology in Alzheimer’s disease

**DOI:** 10.64898/2026.09.26.754651

**Authors:** Agnieszka Urbanek, Alexander J. Dear, David P. Rhodes, Emma F. Garland, Hollie E. Wareing, Harry J. Baker, Willem Molenkamp, Ruixin Zhang, Gefei Chen, Axel Abelein, Simon M. Bell, Juan A. Varela, Thomas C.T. Michaels, J Robin Highley, Georg Meisl, Jan Johansson, Suman De

**Author notes:** Correspondence (G.M.), (J.J), (S.D.). These authors contributed equally (Agnieszka Urbanek, Alexander J. Dear).

## Abstract

Neurotoxic intermediates formed during amyloid-β(Aβ) aggregation are thought to drive Alzheimer’s disease, yet their rarity and transient nature have obscured their identity and formation pathways in the human brain. Here, we combine super-resolution microscopy and single-molecule functional imaging with kinetic modelling to classify Aβ intermediates according to their membrane-disrupting activity and connect their damaging effects to the aggregation pathways that generate them. Applying this framework to recombinant Aβ42, an Alzheimer’s knock-in mouse model and human Alzheimer’s brain tissue, we find two functionally distinct intermediate populations present across all systems. They form through separate aggregation routes and differ in their neurotoxicity, abundance and susceptibility to inhibition. Both damage neurons and glia, but only one converts into plaques. Suppressing these intermediates pharmacologically in mice reduces plaque burden, attenuates astrogliosis and improves cognition. Our findings connect model systems with human pathology and identify intermediate formation pathways as actionable therapeutic targets across protein-misfolding disorders.

## Introduction

Alzheimer’s disease (AD) is the most prevalent form of dementia, affecting over fifty million individuals worldwide. One of its defining pathological hallmarks is the extracellular deposition of amyloid-β (Aβ) peptides into plaques^1^, a process that begins with the misfolding and aggregation of Aβ monomers into soluble intermediates, which subsequently assemble into insoluble fibrils that accumulate as amyloid plaques^2,3^. As Aβ aggregates and deposits in the brain, levels of soluble Aβ in cerebrospinal fluid decline, providing one of the earliest detectable abnormalities in AD^4^. Yet the relevance of this aggregation pathway to disease progression remains contentious: Aβ plaque burden correlates poorly with cognitive decline^5,6^, and immunotherapies aimed at clearance of plaques have shown only modest clinical benefit^7,8^. These observations raise fundamental questions about the causal role of Aβ plaques in AD^9,10^. Mounting evidence instead implicates Aβ intermediates, such as oligomers, formed en route to plaques, as key drivers of neurodegeneration and better correlate with cognitive dysfunction^11–16^.

Among the Aβ isoforms, the 42-residue Aβ peptide, Aβ42, is particularly prone to aggregation^17^ and disproportionately enriched in amyloid plaques in AD brains^18^. It is also deposited into amyloid plaques earlier in the disease process compared to other Aβ isoforms^19^. As a result, the role of Aβ42 in AD is extensively studied in model systems, primarily through recombinant protein assays and transgenic animal models. Whilst these approaches have provided key insights into Aβ dysfunction and associated toxicity, they have certain limitations. Recombinant studies typically require Aβ concentrations several orders of magnitude higher than those found in the human brain^20^. Even the most advanced knock-in mouse models fail to fully recapitulate the spatial and temporal complexity of human AD pathology^21,22^. These inconsistencies, along with the limited success of plaque-targeting therapies, raise doubts about whether aggregation pathways observed in model systems truly parallel those in the human brain. They further cast doubt on whether intermediates detected in controlled assays are present in AD tissue and whether these are indeed the toxic species driving neuronal injury. Without a direct molecular and functional bridge linking model systems to human disease, the identity of the species that initiate the pathogenic cascade, and the pathways by which they form in the brain, have remained elusive.

Progress has been hindered by the fact that Aβ aggregation cannot be monitored directly in the living human brain. Histopathological analyses of postmortem tissue provide static snapshots, the gold standard for confirming the presence of pathology, but offer little insight into the underlying aggregation dynamics. Although structural methods such as optical imaging^23,24^, cryo-EM^25,26^, or NMR^27,28^ have detected Aβ aggregates in brain-derived extracts, they cannot link these structures to their functional consequences or to the pathways that give rise to them. Conversely, functional assays demonstrated the neurotoxicity of brain-derived Aβ assemblies^11,29^, yet they offer no information on how intermediate species emerge or evolve in the diseased brain. Consequently, current therapeutic strategies continue to emphasize late-stage plaque removal, long after irreversible neurodegeneration has occurred, reinforcing the prevailing notion that early intermediates remain effectively undruggable^30^.

To address this challenge, we developed a digital platform that detects individual Aβ species based on their damaging potential at single-aggregate resolution. This assay quantifies Aβ’s ability to disrupt membranes and trigger Ca^2+^ influx. The assay builds on extensive evidence that Aβ toxicity in AD involves disruption of neuronal membranes and Ca^2+^ dyshomeostasis, which in turn lead to synaptic failure, neural network dysfunction, and cognitive decline^31–34^. Aβ-induced Ca^2+^ dysregulation has been directly linked to memory deficits^35^ and Ca^2+^ alterations have been observed in individuals with mild cognitive impairment^36^. Yet the identity of the responsible species, and how to intercept their formation to prevent damage, remain unresolved. Our assay overcomes this limitation by sampling Aβ species formed along the *in vitro* aggregation pathway as well as in mouse models and human postmortem Alzheimer’s brains. It detects Aβ-induced membrane permeabilization and Ca^2+^-ion influx in real time with high sensitivity, registering events even when triggered by the passage of a single Ca^2+^-ion. This enables classification of toxic Aβ species by the strength of their membrane-damaging activity.

Using this approach, we identified two functionally distinct Aβ aggregation intermediates present across recombinant peptides, Alzheimer’s mouse models, and human brain. By integrating functional data with kinetic theory, we mapped how these intermediates form, evolve, drive toxicity, and respond to therapeutic intervention. Our findings establish a link between models and human disease, connect neurotoxic potency to aggregation pathways, and provide a framework for selectively targeting the transient Aβ assemblies that drive neurodegeneration and AD progression.

## Results

### Functional fingerprinting of Aβ aggregates at the single-aggregate level

To quantify the membrane-disrupting activity of individual Aβ aggregates, we developed a single-vesicle assay that reports aggregate-induced damage with single-aggregate resolution. Nanoscale vesicles composed of neuronal membrane-mimicking lipids were loaded with a dextran-conjugated Ca^2+^-sensitive fluorogenic dye (Cal-520) within their lumens. The vesicles were then immobilized in a Ca^2+^-containing buffer on a coverslip (Fig. 1a). In this configuration, any disruption of the vesicle membrane by Aβ aggregates allows external Ca^2+^ to enter the lumen, where it binds to the encapsulated Cal-520 and generates a fluorescence signal that directly reports aggregate-induced damage. We quantified this increase in fluorescence from hundreds of individual vesicles simultaneously using total internal reflection fluorescence (TIRF) microscopy. Because each vesicle functions as an independent reaction chamber with an attoliter-scale (10^-18^ L) volume, and concentrations were chosen such that at most one Aβ aggregate encountered a vesicle at any given time, this design allowed us to quantify the damaging potential of hundreds of aggregates simultaneously with single-aggregate resolution (Fig. 1b).

**Figure 1.**
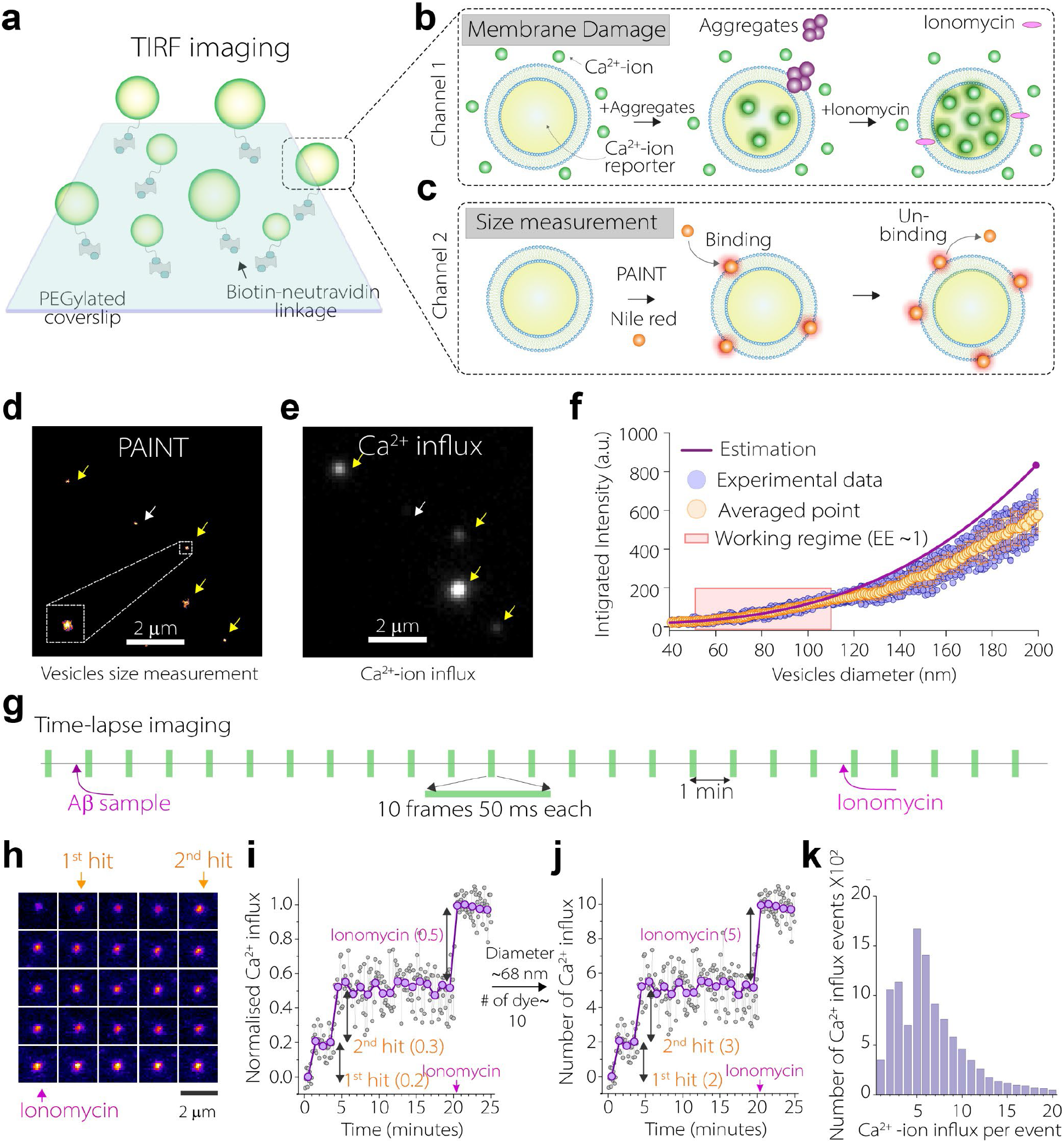
Digital single-vesicle assay for quantifying the number of Ca^2+^ ions per membrane-disrupting hit from individual Aβ aggregates. **(a)** Nanosized lipid vesicles tethered to PEGylated coverslips via biotin-neutravidin linkage and imaged by TIRF microscopy. **(b)** Vesicles loaded with a Ca^2+^-sensitive dye showing increased fluorescence upon Ca^2+^ influx following membrane permeabilization. **(c)** Vesicle size determination using Nile Red-based super-resolution PAINT. **(d, e)** Representative PAINT images of vesicles used for size measurement (d) and vesicles displaying Ca^2+^ influx after ionomycin addition (e). **(f)** Encapsulation efficiency estimated by plotting the fluorescence intensity of individual vesicles (blue circles) against their diameters. Orange circles show the averaged intensity for binned diameter ranges, and the purple line represents the estimated fluorescence expected if the vesicle lumen were completely filled with dye. **(g)** Time-lapse acquisition scheme. Ten frames (50 ms exposure each) were acquired once per minute. Aβ42 was added after the first minute, followed by 20 minutes of imaging and a further 5 minutes after ionomycin addition. **(h)** Montage of a single vesicle showing averaged 10-frame bursts and discrete Ca^2+^ influx events after Aβ42 exposure and ionomycin treatment. **(i-j)** Fluorescence traces from the same vesicle shown in analog (I) and digital (J) modes. Grey circles indicate the intensity of each individual frame within the 10-frame burst, and purple circles represent the averaged intensity of each burst acquired once per minute. A ‘hit’ denotes a detected Ca^2+^-ion influx event in the same vesicle during the time-lapse measurement. **(k)** Distribution of Ca^2+^ influx events (hits) per vesicle following Aβ42 exposure, quantified across hundreds of analyzed vesicles.

Because our aim was to establish a regimen where we could count the exact number of Ca^2+^ ions entering after an Aβ aggregate ‘hits’ the vesicle with single-ion resolution and use this signature as a fingerprint to distinguish different Aβ species, we determined the number of dye molecules encapsulated within each vesicle. We measured vesicle diameter *in situ* using super-resolution PAINT (Point Accumulation for Imaging in Nanoscale Topography) with Nile Red (Fig. 1c). Stochastic binding of Nile Red to vesicle membranes produced ‘blinking’ events that we localized with sub-diffraction precision (Fig. 1d), allowing us to estimate vesicle diameters down to ∼50 nm (Supplementary Fig. 1a). Then we estimated the total number of dye molecules present within each vesicle by multiplying the calculated volume by the known encapsulated Cal-520 dye concentration used during vesicle preparation. To determine the maximal fluorescence signal for each vesicle, we added ionomycin, a Ca^2+^ ionophore, at the end of the experiment to saturate the lumen with Ca^2+^ (Fig. 1e). This ensured that all Cal-520 molecules were bound to Ca^2+^, generating the maximal fluorescence signal and providing a direct correlation between dye number and fluorescence intensity. Cal-520 fluorescence was linear across the working concentration range, confirming the absence of self-quenching artifacts (Supplementary Fig. 1b). The measured fluorescence scaled proportionally with vesicle size, as confirmed by calculating the encapsulation efficiency (EE), the ratio of measured fluorescence intensity to theoretical dye content, for vesicles of different diameters. Consistent with previous reports^37^, the EE was close to unity for vesicles with diameters below 100 nm but declined for larger vesicles (Fig. 1f). Therefore, vesicles between 50 and 100 nm in diameter were selected for subsequent analyses.

We next exposed vesicles to Aβ42 aggregates and monitored for 20 minutes using time-lapse imaging. We observed discrete, stepwise increases in fluorescence, indicating transient bilayer disruptions followed by bursts of Ca^2+^ influx triggered by damaging Aβ42 (Fig. 1g-i). These findings suggest that Aβ42 intermittently compromises neuronal membrane-mimicking bilayer integrity and permits bursts of Ca^2+^ entry, as shown in earlier work^38,39^. By correlating total fluorescence intensity with the number of encapsulated dye molecules, we calculated the number of Ca^2+^ ions entering a vesicle after each hit (Fig. 1j). Each discrete fluorescence step corresponded to a single permeation event that produced a burst of Ca^2+^ entry with single-ion resolution. Moreover, the assay is sensitive enough to detect even an influx of a single Ca^2+^ ion (Supplementary Fig. 2), because the vesicle lumen is on the order of attoliters and the entry of a single Ca^2+^ ion produces a micromolar-scale concentration change that lies well above the dissociation constant (K_d_) of our Ca^2+^ reporter. When we repeated the measurements under low aggregate concentrations, where only ∼10-20% of vesicles encounter aggregates, the Ca^2+^-ion influx distributions remained unchanged, supporting the interpretation that the discrete Ca^2+^ influx signatures arise predominantly from single aggregate-vesicle interactions (Supplementary Fig. 3). The frequency of hits reflects the abundance, whilst the magnitude of influx per hit quantifies the damaging capacity of each species. Together, these parameters provide quantitative measures of the total damaging effect of a sample (Fig. 1k).

### Identification of two classes of functionally distinct Aβ42 intermediates

To track the emergence and evolution of the membrane-disrupting species during aggregation, we applied our assay to recombinant Aβ42 monomers (3 μM) undergoing fibrillization. Aggregation was monitored using Thioflavin T (ThT) fluorescence (Fig. 2a). In parallel, identical aggregation assays were performed without ThT, and aliquots were sampled at defined time points to measure membrane-disrupting activity. In the lag phase of aggregation (e.g., at 15 and 50 minutes), prior to any fibril formation detected by ThT, the Ca^2+^ influx distributions were single-peaked at ∼5 ions per event, defining a population we term class A intermediates (Fig. 2b). As aggregation continued, the frequency of class A events increased and peaked at the midpoint of aggregation (85 min), and declined thereafter, indicating that these class A species are transient and likely consumed or transformed during fibril elongation. By 480 minutes, the influx distribution again became single-peaked, centered at 2-3 Ca^2+^ ions per event (Fig. 2b). These relatively less-damaging species, designated as class B, first emerged at the end of the lag phase. Their abundance peaked at ∼150 minutes, and gradually decreased thereafter. Despite this decline, class B aggregates remained detectable throughout the remainder of the aggregation process. In addition to these dominant A and B species, we observed a distinct influx distribution at the very early stages of aggregation (3 minutes), characterized by its low-magnitude signals with a peak at one ion per event and a fast decay. We refer to this rare, early-forming, and weakly damaging distribution as class C. Each class is defined by its characteristic distribution of Ca^2+^ influx events, and the peak positions (5, ∼2-3, and 1 ions per event for Class A, Class B and Class C, respectively) are used only as convenient descriptors throughout the manuscript.

**Figure 2.**
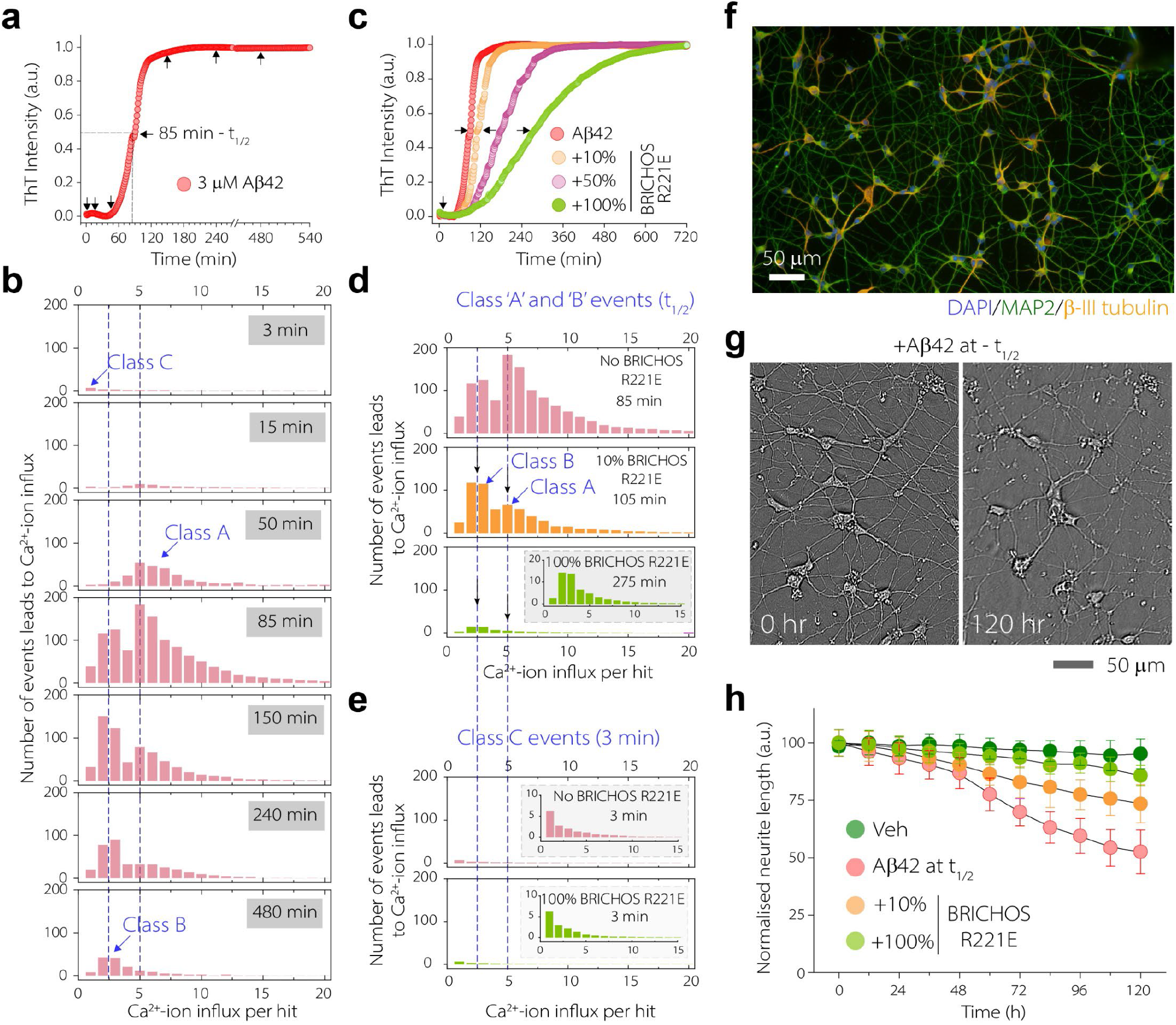
Two types of intermediates emerge during Aβ aggregation. **(a)** Aggregation of 3 μM recombinant Aβ42 monitored using ThT fluorescence, with arrows marking the specific time points chosen for subsequent analyses. **(b)** Ca^2+^-ion influx distributions at indicated time points. **(c)** Aggregation kinetics of Aβ42 in the presence and absence of BRICHOS R221E monomers of 10%, 50%, or 100% molar equivalents relative to Aβ42 monomer, with selected time points for analysis indicated by arrows. **(d-e)** Ca^2+^ influx distributions from Aβ42 samples collected at the aggregation half-time (t1/2) determined for each individual aggregation trace (d) or at 3 min (e), with (10% or 100%) and without BRICHOS R221E treatment. **(f)** Immunocytochemistry of hPSC cortical neurons stained with DAPI (blue), MAP2 (green), and β-III-tubulin (orange) (n = 4 independent differentiation). **(g-h)** Live imaging of hPSC neurons exposed to 1 µM Aβ42 aggregates collected at the aggregation half-time, prepared either in the presence or absence of BRICHOS, with PBS as the vehicle control. Representative images after Aβ42 exposure prepared without BRICHOS at 0 h and 120 h (g), and quantification of neurite length over 5 days for all conditions (h). Data points are plotted as the mean ± standard deviation of three biological replicates.

These class C species were present at low levels and did not significantly contribute to membrane activity at later time points. To confirm the robustness of these classifications, we repeated Aβ42 aggregation at different concentrations (2 and 5 μM) and observed similar patterns of Ca^2+^-ion influx peaks (Supplementary Fig. 4a-d) corresponding to class A and class B intermediates. We also plotted the number of events at each Ca^2+^ influx value over time normalized to 1 (Supplementary Fig.. 5). The resulting curves for low and high Ca^2+^ influx values clearly revealed two distinct kinetic behaviours corresponding to the rise and decline of our proposed class A and B species over the course of aggregation, further justifying this classification approach.

To probe the underlying pathways that lead to the formation of these toxic species, we aggregated Aβ42 in the presence of varying concentrations of BRICHOS R221E monomer, a chaperone known to slow down aggregation by specifically inhibiting secondary nucleation^40,41^ (Fig. 2c). For each condition, samples were collected at the half-time (t_1/2_) of the corresponding aggregation trace. We used t_1/2_ because it provides a reproducible reference point at which fibril mass reaches ∼50% of its final level, enabling comparison of aggregate populations at an equivalent mechanistic stage of fibril formation despite differences in aggregation rate. The addition of BRICHOS at the start of the aggregation led to dose-dependent suppression of both class A ( peaked at5 Ca^2+^ ions) and class B (peaked at ∼2-3 Ca^2+^ ions) distribution (Fig. 2d). At 10% molar equivalents relative to monomeric Aβ42, BRICHOS reduced the class A peak by ∼75%, and at 100% BRICHOS, it was completely eliminated. In contrast, class B events remained detectable even at the highest BRICHOS level (100%) (Fig. 2D). Because this assay detects only membrane-disruptive intermediates and not nondamaging ones, the reduction in events reflects a selective loss of class A and class B species or a loss of their damaging capacity in the presence of BRICHOS, rather than implying that intermediates dissociate or that aggregation proceeds without intermediate formation. By comparison, class C events were unaffected, even at 100% BRICHOS concentration (Fig. 2e). This behaviour cannot be attributed to a direct effect of BRICHOS on vesicle integrity, as this chaperone alone does not induce Ca^2+^ influx in the absence of Aβ (Supplementary Fig. 6). Although the membrane-mimicking vesicles used here do not fully capture native membrane complexity, their controlled composition enables quantitative single-aggregate comparison of membrane-disrupting activity. Future studies using native membranes will be needed to determine how physiological membrane composition modulates the function of each intermediate class.

We then asked whether this BRICHOS R221E-mediated suppression of membrane-disrupting class A and B intermediates translates into neuroprotection. To test this, human pluripotent stem cell (hPSC)-derived cortical neurons (Fig. 2f) were exposed to Aβ42 aggregates collected at the midpoint of aggregation, when both class A and class B species are present. Aggregates were prepared either in the absence or presence of BRICHOS R221E, and neuronal connectivity was measured by quantifying neurite length every 6 hours over five days using live-neuron imaging (Fig. 2g). This neurite connectivity assay parallels disruption of long-term potentiation, a well-established cellular correlate of learning and memory^42^. Aβ42 intermediates caused progressive neurite shortening, whereas BRICHOS R221E treatment conferred dose-dependent protection (Fig. 2h). Together, these results establish that Aβ42 aggregation generates distinct classes of neurotoxic intermediates with characteristic damaging profiles, and that their targeted suppression provides direct protection to human neurons.

### Distinct Aβ42 intermediates have distinct kinetic signatures and damaging potential

At each stage of Aβ42 aggregation, the Ca^2+^ influx distributions showed peaks at ∼2–3 ions, ∼5 ions, or both, corresponding to class B and class A species with distinct kinetic behaviours. To quantify these species accurately, we first defined pure fingerprints for each class. For class A, we selected the end of the lag phase (50 minutes for 3 μM Aβ42), when the influx distribution showed only the ∼5 ion peak. This profile also closely resembled that at 15 minutes, early in the lag phase but without the small contributions from the rare class C species (Fig. 3a). For class B, we used the aggregation half-time in the presence of 100% BRICHOS R221E, a condition that completely suppressed class A activity and thus yielded a pure class B profile. This distribution also corresponds to the end of aggregation, when only class B species remained (Fig. 3b).

**Figure 3.**
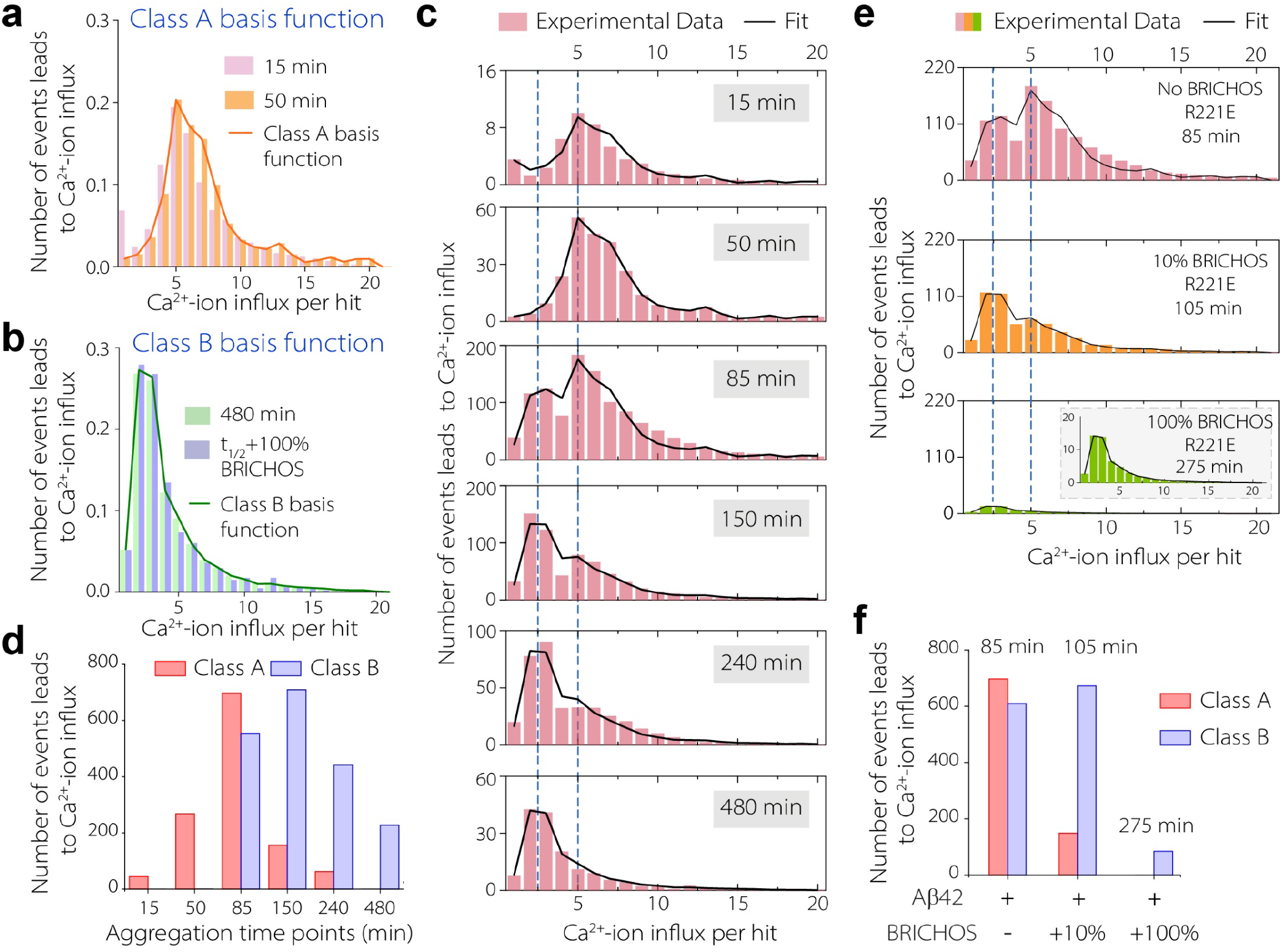
Distinct kinetic profiles of membrane-disrupting Aβ42 aggregates. **(a-b)** Basis distributions used to define class A and class B intermediates, derived from Ca^2+^ influx profiles. Class A corresponds to the distribution observed during the lag phase (a), and class B corresponds to the distribution at the end of aggregation (b). **(c)** Experimental Ca^2+^ influx distributions at selected time points with representative fit overlays for 3 µM Aβ42. **(d)** Quantified abundance of class A and class B intermediates across aggregation time points. **(e)** Ca^2+^ influx distributions of aggregates collected at the aggregation half-time and prepared with varying concentrations of BRICHOS R221E or without, with fit overlays. **(f)** Quantified contributions of class A and class B intermediates in aggregation half-time samples prepared with or without BRICHOS R221E.

We then fitted the Ca^2+^ influx distributions at each aggregation time point to a linear superposition of these fingerprint distributions. We found that, at every stage, membrane activity could be explained by either one species alone or by a mixture of the two, and the overall distributions can be described as linear combinations of their contributions (Fig. 3c), accounting for the full spectrum of Aβ42-induced membrane-disrupting activity. From this fitting, we quantified the contribution of class A and class B events at different aggregation time points, estimated their relative abundance and tracked their evolution across the aggregation trajectory (Fig. 3d). Class B aggregates followed a classical oligomerization pattern: their abundance increased after the lag phase, peaked during the growth phase, and declined once fibril formation saturated. In contrast, class A aggregates appeared earlier, peaking before substantial fibril formation. This timing is inconsistent with a pathway that follows fibrillization and instead indicates that class A species originate through a distinct kinetic route.

Finally, we fitted the Ca^2+^ influx distributions obtained in the presence of BRICHOS using the same fingerprint superposition to examine how it modulated the relative abundance of the two classes (**Fig. 3e**). The fits showed strong differential sensitivity: class A was completely suppressed by BRICHOS, whereas class B remained detectable even at the highest concentration tested (**Fig. 3f**). This differential response further supports that class A and class B arise through distinct aggregation pathways.

To further probe whether these functional classes correspond to physically distinct assemblies, we performed super-resolution imaging of aggregates collected at 50 and 180 minutes, which revealed that Class A aggregates are smaller on average than Class B (Supplementary Fig. 7a-d). We also analyzed samples collected at 85 minutes, when both classes are present, using sucrose gradient fractionation. In these experiments, lower-density fractions were enriched in class A species, whereas higher-density fractions were enriched in class B aggregates (Supplementary Fig. 7e-g). ANS fluorescence analysis further showed a blue-shifted emission spectrum in the Class A-enriched fraction, suggesting greater exposed hydrophobicity (Supplementary Fig. 7h). Together, these data suggest that Class A and Class B are not only functionally distinct, but also differ in size and surface hydrophobicity.

### Global kinetic analysis reveals distinct origins of intermediates

We next sought to contextualize the emergence of class A and class B species within the broader kinetics of Aβ42 aggregation. To this end, we performed global kinetic fitting of Aβ42 fibril formation data alongside the extracted kinetic curves for both species across multiple monomer concentrations (Fig. 4a). We first applied a state-of-the-art model in which all intermediates are predominantly formed via secondary nucleation (Fig. 4b), consistent with canonical Aβ42 oligomerization mechanisms and previously reported rate constants^43,44^. This model failed to capture the early rise in class A species across concentrations (Fig. 4c) but adequately described the kinetics of class B species (Fig. 4d). We therefore tested an alternative model in which a much larger proportion of class A species are formed via primary rather than secondary nucleation, in contrast to what is observed for fibrils^43^ or directly measured oligomers^44^. This model produced substantially better agreement with the experimental data, particularly at early time points when class A species are most abundant.

**Figure 4.**
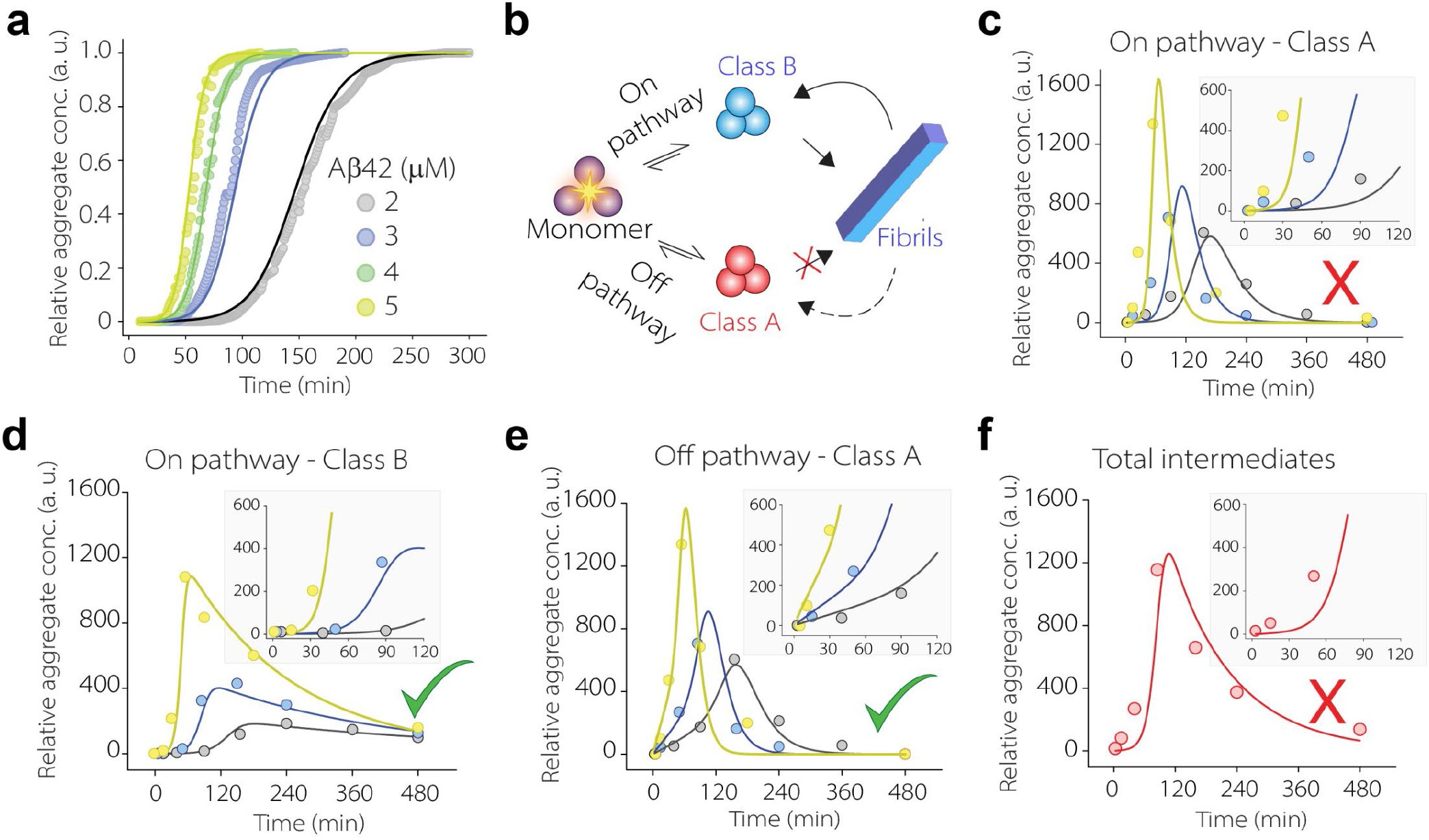
Global kinetic fitting shows distinct origins of class A and class B intermediates. **(a)** Aggregation kinetics of Aβ42 monitored by ThT fluorescence (points), with global kinetic fits overlaid (lines). **(b)** Schematic representation of on-pathway and off-pathway routes for formation of intermediate aggregates. **(c-d)** Global fits of class A (c) and class B (d) intermediates, using a model in which both species are generated via secondary nucleation, only through the on-pathway route. **(e)** Alternative model incorporating at least partial off-pathway formation of class A species through primary nucleation or fibril-independent mechanisms. **(f)** Combined class A and class B intermediates fitted under the on-pathway model, which does not capture the combined intermediate profile Insets (c-f) show early aggregation stages.

Together, these analyses indicate that class A and class B species arise through distinct kinetic routes, and have important implications for their connection to fibrils. Since fibrils are formed by primary and secondary nucleation in the same relative proportions as class B species, this implies that they are likely on-pathway intermediates of fibril nucleation and can be identified with the oligomeric intermediate species detected previously^44^. On-pathway in this context means that a substantial fraction of the overall nucleation flux passes through these intermediates^45^. It does not mean that most class B oligomers ultimately become fibrils; like all amyloid oligomers, both on- and off-pathway species are typically transient and predominantly dissociate back into monomers^45,46^. By contrast, class A species can contribute only minimally to fibril formation; if they did, a much larger share of fibrils would arise via primary nucleation. Consistent with this, we found that BRICHOS strongly inhibits class A species, and yet has no detectable impact on fibril primary nucleation^47^. This implies that class A species is off-pathway, contributing little to fibril formation^45^ (Fig. 4e).

When both populations were combined and treated as a single intermediate pool, as in earlier studies^44^, the model underestimated aggregate formation during the lag phase (Fig. 4f). Although this by itself does not prove the existence of two discrete intermediates, it exposes a fundamental inconsistency with a purely on-pathway mechanism.

### Alzheimer’s brain contains the same two classes of intermediates

To assess the disease relevance of the intermediates of Aβ42 aggregation identified *in vitro*, we next asked whether class A and class B species are also present in the human brain. We examined postmortem (PM) frontal cortex tissue from three sporadic AD patients. In each case, we confirmed plaque pathology by immunohistochemistry using Aβ40-specific antibody (EPR23712-2, Abcam) and Aβ42-specific antibody (21F12, Absolute antibody) (Fig. 5a, Supplementary Fig. 8). We found that more than 90% of amyloid plaques are composed of only Aβ42. Then, to isolate diffusible intermediates, we applied a gentle extraction protocol previously shown to release diffusible, neurotoxic Aβ species^48,49^. Both small Aβ fibrils^25^ and neurotoxic oligomers^42,48^ have been detected in this fraction. However, mature fibrils have been shown to contribute minimally to membrane damage^50^. Therefore, any membrane-disruptive activity detected in our assay is expected to reflect diffusible Aβ intermediates.

**Figure 5.**
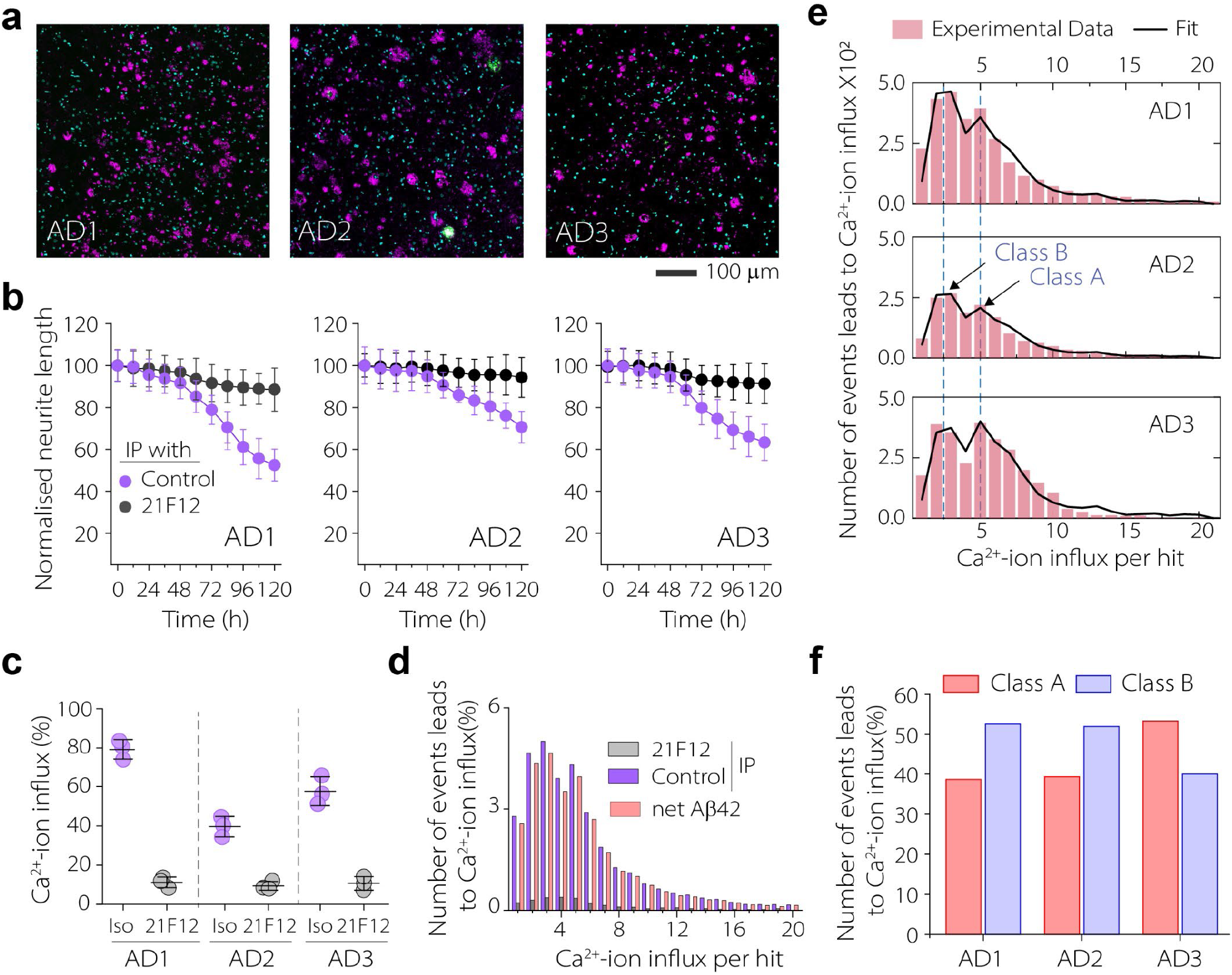
Human Alzheimer’s brain contains two classes of toxic Aβ42 intermediates as observed in the recombinant system. **(a)** Immunohistochemistry of frontal cortex from three AD cases, stained with the Aβ40-specific antibody EPR23712-2 (green), Aβ42-specific antibody 21F12 (purple), and DAPI (cyan). **(b)** Time-course of neurite outgrowth in hPSC-derived cortical neurons treated with immunoprecipitated (IP) brain extracts using either 21F12 or an isotype control antibody. **(c)** Quantification of Ca^2+^ influx activity for the same IP samples. Data points are plotted as the mean ± standard deviation of three biological replicates(b, c) **(d)** Net Aβ42-induced Ca^2+^ influx distributions from brain extracts (example shown: AD1), calculated by subtracting 21F12-IP from isotype control. **(e)** Net Ca^2+^ influx distributions fitted using class A and class B basis functions. **(f)** Relative contributions of class A and class B intermediates for each AD case.

To determine the specific contribution of Aβ42, brain extracts were immunoprecipitated with the Aβ42-specific 21F12 antibody or an isotype control. Fractions depleted with the control antibody showed higher levels of Aβ42 aggregates than those treated with 21F12, as confirmed by single-aggregate imaging (Supplementary Fig. 9). Functionally, these control-antibody depleted fractions also induced pronounced neurite degeneration in hPSC cortical neurons (Fig. 5b) and showed strong membrane-disrupting activity (Fig. 5c). In contrast, fractions immunodepleted with 21F12 displayed lower toxicity in both readouts.

We next quantified the net contribution of Aβ42 by calculating the difference between control and 21F12-antibody depleted fractions (Fig. 5d). The resulting distributions revealed peaks at ∼2-3 and ∼5 Ca^2+^ influx events, matching the class A and class B species identified in a recombinant systems. Fitting these data to their established fingerprint distributions confirmed the presence of both class A and B intermediates in all three AD patients (Fig. 5e), although their relative abundances varied across cases (Fig. 5f). These findings demonstrate that the two neurotoxic classes defined *in vitro* are also found in AD brain, indicating the disease relevance of the intermediates captured by recombinant Aβ peptide.

Although these species are transient in *in vitro* aggregation reactions, their detection in PM brain tissue is not unexpected. Flash-frozen PM tissue captures a snapshot of the *in vivo* aggregation landscape, where continuous Aβ monomer production and ongoing assembly maintain a dynamic reservoir of transient intermediates. By contrast, in vitro reactions lack sustained monomer supply; once monomers are consumed, intermediates diminish as aggregation proceeds to completion.

### *In vivo* suppression of intermediates correlates with therapeutic benefit

To explore whether suppression of Aβ42 intermediates in vivo contributes to therapeutic benefit, we turned to a well-characterized mouse model of AD. The *APP*^*NL-G-F*^ knock-in mouse model expresses humanized Aβ carrying the Arctic (E22G) mutation and shows progressive amyloid pathology, neuroinflammatory response of glial cells and behavioural abnormalities^22,51^. Previous studies have shown that intravenous administration of BRICHOS R221E monomer in this mouse model reduces plaque burden, attenuates astrogliosis, and improves cognition^52,53^. However, the molecular basis of these benefits remains unclear. Using tissue from the same study, we confirmed reductions in plaque load (Fig. 6a) and astrogliosis (Fig. 6b) and asked whether these therapeutic benefits are linked to a suppression of Aβ intermediates identified in our *in vitro* experiments.

**Figure 6.**
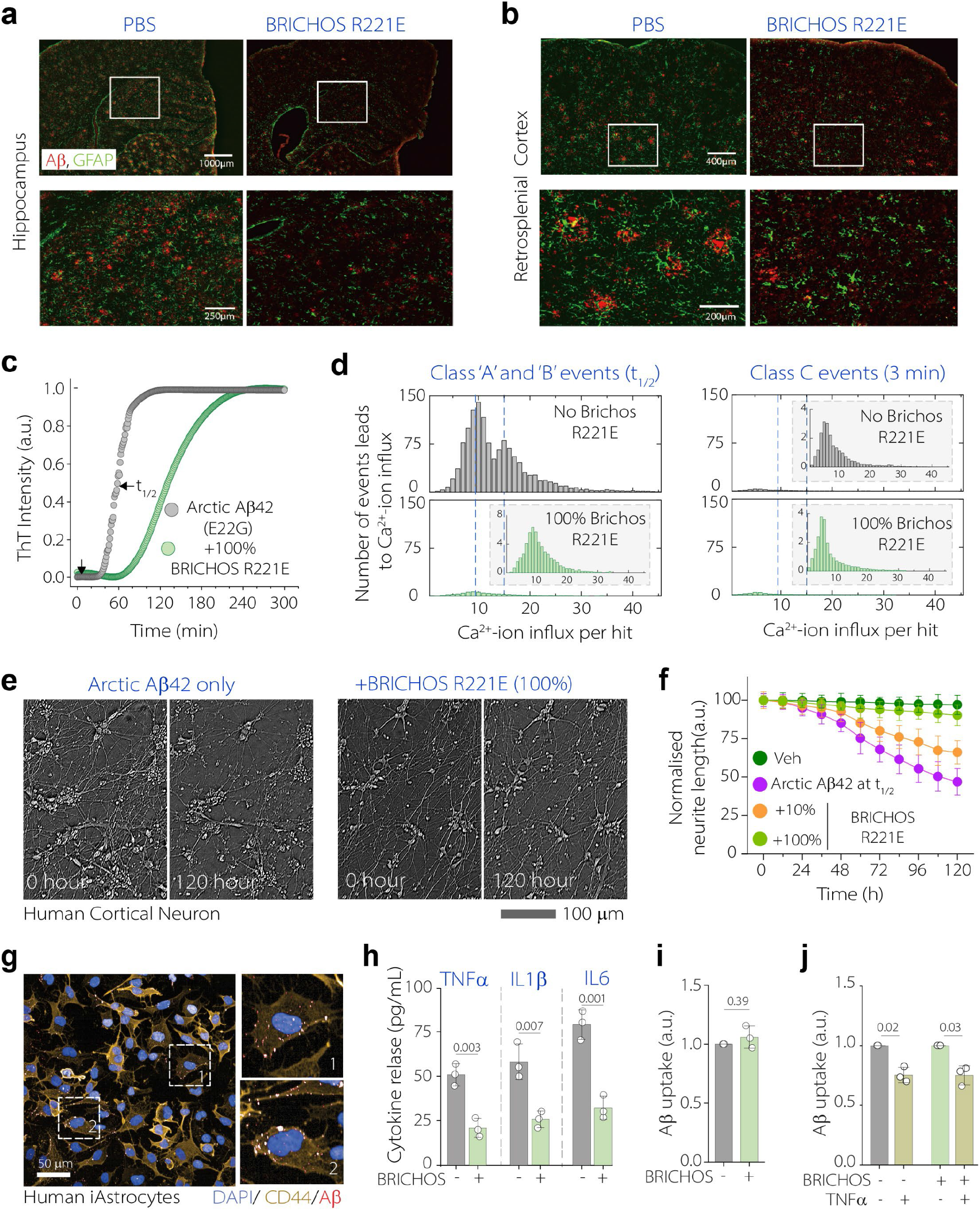
BRICHOS R221E suppresses the formation Arctic Aβ42 oligomeric intermediates and attenuates neuronal and astrocytic dysfunction. **(a-b) I**mmunohistochemistry images of hippocampus (a) and retrosplenial cortex (b) from PBS-, and BRICHOS R221E–treated *APP*^*NL-G-F*^ mice stained for astrocytes (GFAP, green) and Aβ (82E1, red) (n = 4 mice/group). **(c)** Aggregation kinetics of 3 μM Arctic Aβ42 in the absence or presence of 100% BRICHOS R221E. **(d)** Ca^2+^ influx distributions induced by Arctic Aβ42 at the aggregation midpoint and at the early time point ( 3 min) with or without 100% BRICHOS R221E. **(e)** hPSC neurons exposed to Arctic Aβ42 at t1/2 prepared in the absence or presence of 100% BRICHOS R221E. **(f)** Quantification of mean neurite length upon Arctic Aβ42 exposure. **(g)** Immunocytochemistry images of human iAstrocytes treated with Arctic Aβ42 aggregates for 1 hour. **(h)** Quantification of secreted proinflammatory cytokines in iAstrocyte culture medium in response to Arctic Aβ42 aggregates prepared with or without 100% BRICHOS R221E. **(i)** Uptake of Arctic Aβ42 aggregates prepared in the absence or presence of BRICHOS, measured by 6E10 antibody. **(j)** Uptake of Arctic Aβ42 aggregates by astrocytes pre-incubated with TNF-α for 24 h. Data points are plotted as the mean ± standard deviation of three biological replicates (f. h-j)

As a first step, we aggregated recombinant Arctic Aβ42 (Fig. 6c) and characterized intermediates formed in the aggregation pathway. Although Arctic intermediates caused stronger membrane disruption than wild-type, consistent with the reported higher membrane-disrupting potency of Arctic Aβ42 attributed to increased transmembrane stability^54^, the overall pattern remained similar. At the midpoint of aggregation, we observed two peaks of Ca^2+^ influx similar to class A and class B species. The addition of BRICHOS R221E suppressed both classes of intermediates, with a stronger effect on class A species (Fig. 6d), mirroring its impact on wild-type Aβ42. Thus, despite causing more membrane damage, Arctic Aβ42 displays the same pattern of Ca^2+^ influx and similar sensitivity to BRICHOS.

We next asked whether these intermediates contribute to the attenuated pathology observed in *APP*^*NL-G-F*^ mice when treated with BRICHOS R221E^52,53^. We collected Arctic Aβ42 aggregates from the aggregation midpoint (t_1/2_) and applied them separately to hPSC neurons or to iAstrocytes. In neurons, Arctic Aβ42 aggregates caused pronounced neurite shortening, which was substantially reduced when aggregates were prepared in presence of BRICHOS R221E (Fig. 6e,f), consistent with the trend observed for wild-type Aβ42. A similar pattern was observed in iAstrocytes: Arctic Aβ42 induced robust secretion of the proinflammatory cytokines TNF-α, IL-1β, and IL-6, whereas aggregates prepared in presence of BRICHOS elicited markedly lower cytokine release (Fig. 6g,h). Because astrocytic uptake is a major route of Aβ aggregate clearance^55^, we next examined whether BRICHOS affects this process. Surprisingly, uptake of Arctic Aβ42, quantified using an Aβ-specific 6E10 antibody, was comparable regardless of whether aggregates were formed with or without BRICHOS (Fig. 6i). However, when iAstrocytes were pre-treated with TNF-α for 24 h to mimic chronic inflammatory activation, uptake was significantly reduced (Fig. 6j). These results indicate that Aβ42 intermediates drive cytokine secretion and create a sustained pro-inflammatory environment that, in turn, impairs subsequent Aβ uptake. Thus, our results suggest BRICHOS reduces plaque pathology not by directly enhancing astrocytic uptake or producing Aβ42 aggregates that are inherently easier to internalize, but by preventing the formation of toxic intermediates that trigger pro-inflammatory signalling and ultimately hinder clearance.

To define how neurotoxic Arctic Aβ42 intermediates evolve, we first measured how Ca^2+^ influx distributions change across the aggregation time course in recombinant system (Fig. 7a). Two distinct populations again emerged: an early, high-influx distribution peaks at ∼15 Ca^2+^ ions (Class A) and a later, lower-influx distribution with a peak at ∼9 Ca^2+^ ions that dominated as fibrils began to form (Class B). As with wild-type, we extracted Arctic Aβ42-specific Ca^2+^ influx fingerprint distributions for class A and B species using the influx profile from the lag phase and from the fibril half-time in the presence of BRICHOS R221E, respectively (Fig. 7b). These two fingerprints were sufficient to reconstruct the Ca^2+^ influx distributions at all aggregation time points as linear combinations of Class A and Class B contributions.

**Figure 7.**
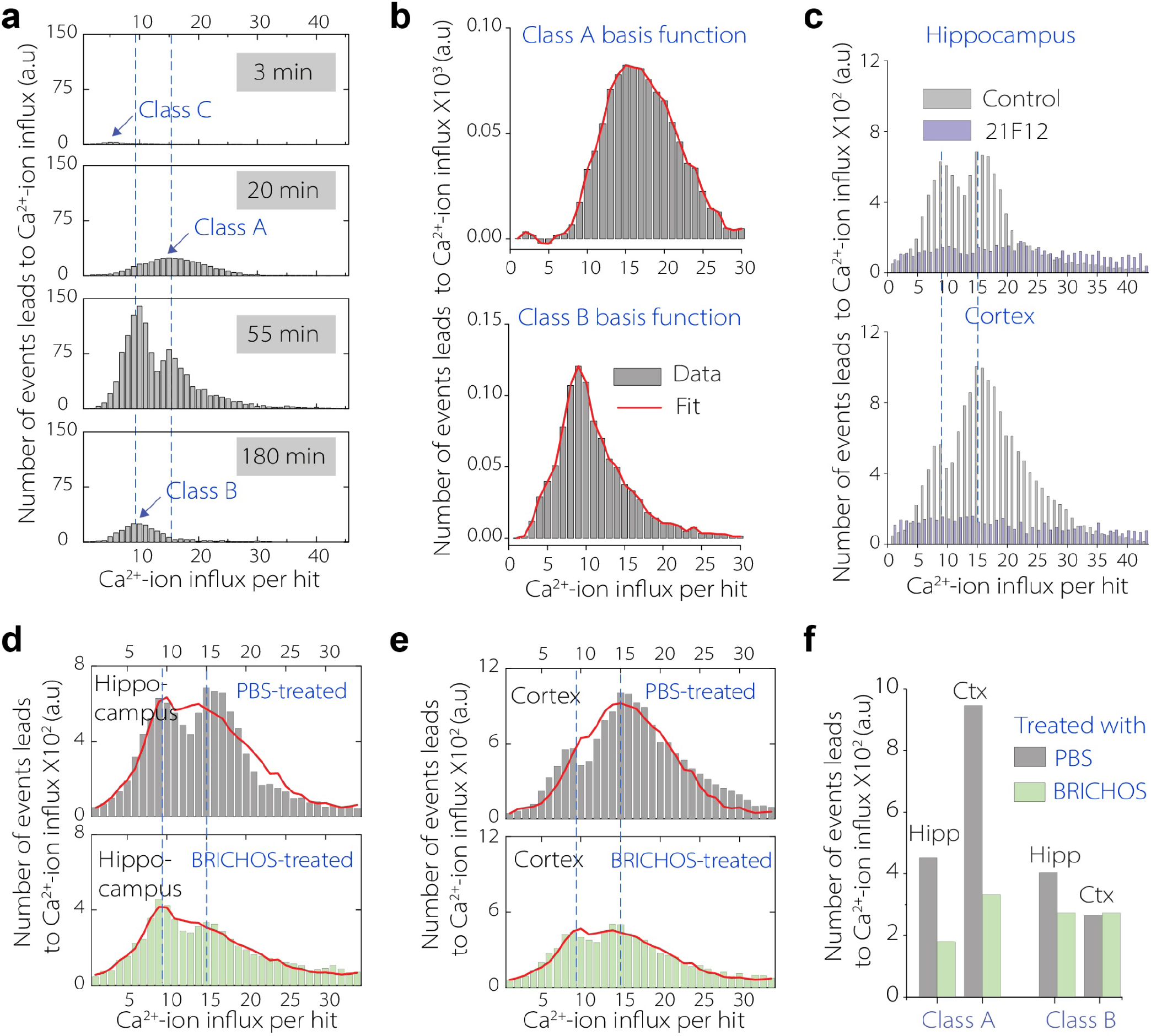
Arctic Aβ42 intermediates identified *in vitro* are also detected in *APP*^*NL-G-F*^ mouse brain and are markedly reduced in BRICHOS-treated mice compared with controls. (**a)** Ca^2+^ influx distributions from Arctic Aβ42 samples collected at the indicated aggregation time points. **(b)** Basis functions for class A and class B species derived from Arctic Aβ42 aggregation at 20 min and BRICHOS-treated t1/2 samples, respectively. **(c)** Ca^2+^ influx distributions from hippocampal and cortical brain homogenates of PBS-treated *APP*^*NL-G-F*^ mice after immunoprecipitation with 21F12 or isotype-control antibodies. **(d-e)** Ca^2+^ influx distributions from hippocampal (D) and cortical (E) brain homogenates of PBS-and BRICHOS R221E–treated *APP*^*NL-G-F*^ mice. **(f)** Estimated proportion of class A and class B events in hippocampus and cortex of PBS- and BRICHOS-treated mice.

We next asked whether these intermediates are suppressed in BRICHOS R221E–treated mice. Homogenates were prepared from the cortex and hippocampus of 12-month-old *APP*^*NL-G-F*^ mice treated intravenously with either BRICHOS R221E or PBS (n = 6 per group). Single-aggregate imaging revealed a significant reduction in diffusible aggregates in BRICHOS-treated mice compared to PBS controls (Supplementary Fig. 10a-f). In PBS-treated animals, both brain regions showed robust Ca^2+^ influx activity, which was significantly reduced after immunoprecipitation with the Aβ42-specific antibody 21F12, confirming that the activity was mediated by Arctic Aβ42 (Fig. 7c). Consistent with this reduction in Aβ42 aggregate abundance after BRICHOS treatment, membrane permeabilization assays revealed a marked decrease in membrane-disrupting activity (Fig. 7d-e, Supplementary Fig.10g). Quantification showed that BRICHOS treatment markedly lowered the number of Ca^2+^ influx events in both hippocampal homogenates (decreasing from 9942 in PBS-treated samples to 6528 in BRICHOS-treated samples) and cortical homogenates (decreasing from 14810 to 8357) (Fig. 7d-e). Similar to our observations from the recombinant system, two distinct Ca^2+^ influx peaks corresponding to Class A and Class B intermediates were detected in brain homogenates from both PBS- and BRICHOS R221E-treated mice across hippocampus (Done 7d) and cortex (Fig. 7e). Fitting these brain-derived distributions with the Class A and Class B basis functions showed that both species were present in PBS-treated mice, whereas BRICHOS R221E treatment reduced the abundance of both classes (Fig. 7f). These results further demonstrate that both classes of Aβ42 intermediates are present *in vivo* and indicate a molecular link between suppression of both intermediate species and the observed therapeutic effects.

## Discussion

In this work, we demonstrate two mechanistically distinct Aβ42 aggregation routes that generate transient, neurotoxic intermediates with different damaging potential. Using a digital single-molecule assay, we defined these two classes of species across recombinant systems, an AD knock-in mouse model, and human Alzheimer’s brain. One class arises early during the lag phase, displays strong membrane-disrupting potency, and appears off-pathway relative to fibril formation. The other emerges later, more closely aligned with fibril precursors, and exhibits lower acute toxicity, but accumulates in greater numbers and thereby contributes substantially to the overall toxic burden. Suppression of both classes *in vivo* reduced plaque deposition, neuronal injury, and gliosis. These findings establish the physiological relevance of distinct aggregation routes and provide a mechanistic framework for understanding how early-stage Aβ intermediates drive AD progression.

*In vitro* studies of recombinant Aβ have revealed a wide spectrum of aggregates that differ in size (dimers, trimers, hexamers), structure (parallel/antiparallel β-sheets, β-hairpin assemblies), and morphology (globular and annular oligomers) formed en route to fibril formation^56,57^. Complementary evidence from human studies also indicates that Aβ intermediates with diverse morphologies exist *in vivo* and act as potent drivers of neurodegeneration^24,58,59^. Yet these species remain difficult to characterize because they are rare, transient, and heterogeneous^60^. As a result, several key questions remain unresolved: Are the intermediates observed *in vitro* the same species that are present in the human brain? Do they follow similar formation pathways *in vivo*? Is their mode of formation directly coupled to specific pathological functions? If so, can these insights be harnessed to design therapies that selectively neutralize pathogenic intermediates at the early stages of disease, when intervention is likely to be most effective?

Our work addresses these questions by introducing a single-molecule assay that integrates super-resolution imaging with functional readouts to classify Aβ assemblies at the single species level by their ability to disrupt membranes and drive calcium dyshomeostasis. Since we observe Ca^2+^ influx as transient events, this suggests that Aβ intermediates induce short-lived membrane permeabilization, potentially through temporary pore formation or local membrane damage. These disruption processes represent key pathways through which Aβ contributes to synaptic failure and neuronal loss in Alzheimer’s pathology^31–36^. Our approach enables detection of rare and transient intermediates, disentangles coexisting species with distinct functions that remain hidden in ensemble measurements, and, when integrated with kinetic modeling, directly links formation pathways to functional outcomes. Such precision is essential because rare but highly toxic assemblies may arise through specific aggregation routes and exert a disproportionate impact on disease progression. Notably, this assay is not restricted to Aβ or to a specific lipid composition, and its modular design makes it broadly adaptable to other misfolding proteins involved in a wide range of protein-aggregation disorders.

Applying our framework to recombinant Aβ42 aggregation revealed a bifurcated kinetic trajectory defined by two mechanistically and functionally distinct intermediates (Fig. 8). Class A species emerge transiently during the lag phase, coinciding with primary nucleation or early oligomeric association steps, and display potent membrane-disrupting activity. Their transient nature reflects intrinsic molecular properties rather than abundance, and their short-lived nature likely explains why they have been largely missed in bulk measurements of soluble oligomers^44^. Class B intermediates, by contrast, accumulate during fibril growth and decline as fibril formation plateaus. Their temporal profile and sensitivity to fibril-surface inhibition are consistent with species generated through secondary nucleation, in line with classical models of fibril-catalyzed oligomer formation and direct oligomer measurements^44^. Furthermore, inhibition of class B species by BRICHOS R221E closely mirrored its suppression of fibril secondary nucleation, which is achieved by binding to and blocking catalytic sites on fibril surfaces^61,62^. Although BRICHOS R221E changed the timing and magnitude of Aβ42 aggregation, the temporal profile of membrane-damaging species was preserved across conditions, with activity rising during early aggregation, peaking around the aggregation midpoint, and declining during the plateau phase (Supplementary Fig. 11). Supporting this, when we bypassed primary nucleation by seeding Aβ42 aggregation with pre-formed fibrils^63^, the lag phase was largely abolished and the reaction generated predominantly Class B species, with minimal Class A detection (Supplementary Fig. 12).

**Figure 8.**
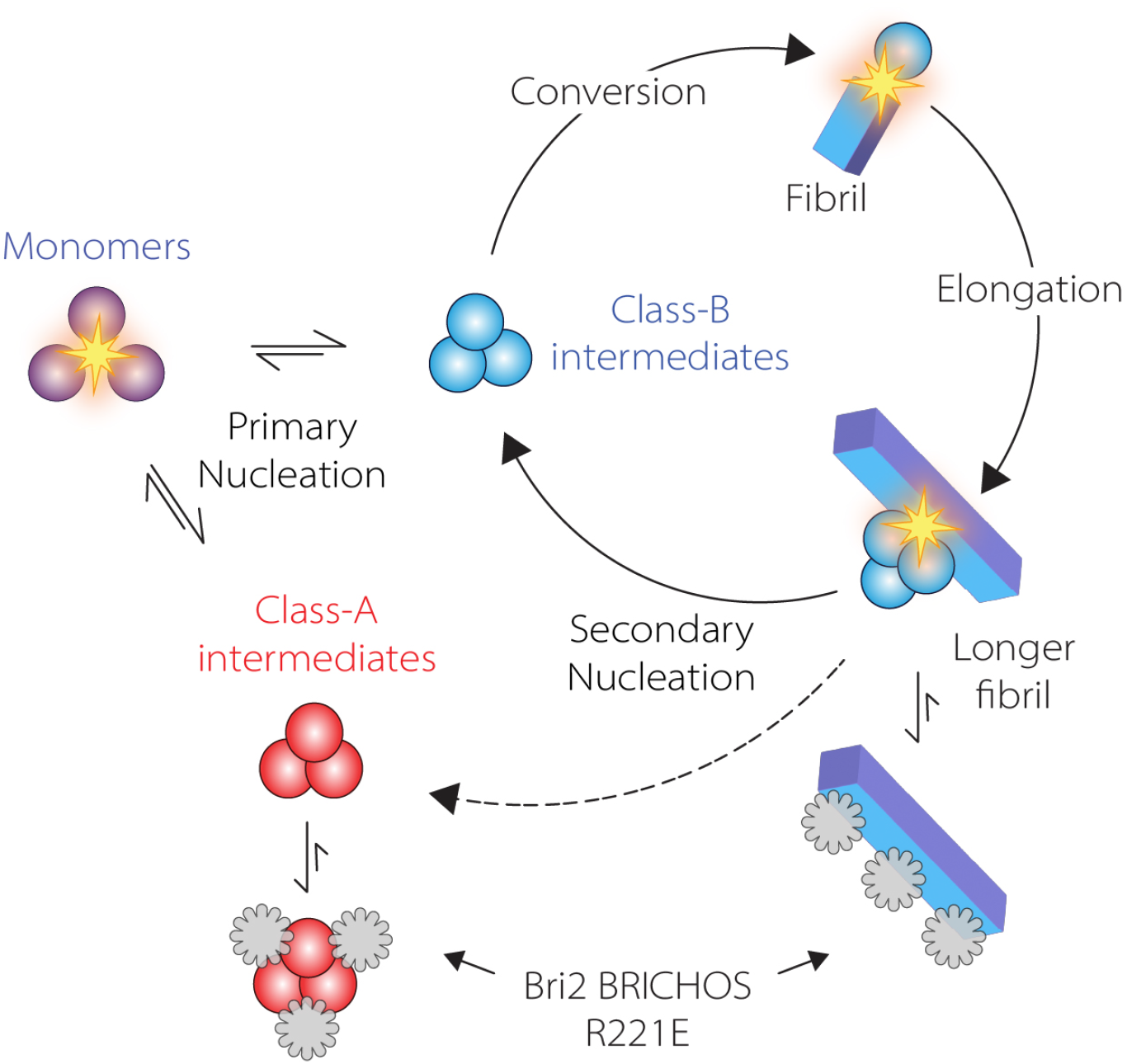
Proposed model of Aβ42 aggregation pathways that generate distinct toxic intermediates. Recombinant Aβ42 undergoes primary nucleation to generate two classes of intermediates. Class A intermediates (red) arise predominantly from primary nucleation. Class B intermediates (blue) emerge during fibril growth, largely through secondary nucleation on fibril surfaces, and can convert into fibrils. BRICHOS reduces Class B formation by binding to fibril surfaces and blocking secondary nucleation, whereas its effect on Class A primarily reflects suppression of the membrane-damaging activity of pre-formed species.

In contrast, Class A intermediates were reduced more strongly than expected from secondary nucleation suppression alone. These species begin to appear during the lag phase, when primary nucleation dominates, suggesting that their formation is not predominantly driven by fibril-surface-catalyzed secondary nucleation. Consistent with this, the small molecule M1 previously shown to suppress Aβ42 secondary nucleation, selectively reduced Class B species, whereas Class A species remained the dominant membrane-damaging population (Supplementary Fig. 13). Moreover, when pre-formed aggregates collected at a time point containing both Class A and Class B species were incubated with BRICHOS R221E, the membrane-disrupting activity of Class A, but not Class B, was strongly reduced (Supplementary Fig. 14). These findings suggest that BRICHOS attenuates Class A activity after formation, likely through direct binding to these off-pathway intermediates, as reported previously^64,65^, rather than by preventing their initial generation. Together, these data support a model in which Class A and Class B species arise through distinct pathways: Class A represents an off-pathway, soluble membrane-active intermediate, whereas Class B is more closely linked to fibril-dependent secondary nucleation.

Distinct Aβ aggregates with different membrane-disrupting activity have been described previously and attributed to differences in hydrophobicity and epitope exposure^20,24^. Previous studies have also identified intermediate populations, some able to convert into fibrils and others that remain fibril-incompetent^66,67^, but the mechanistic basis of this distinction has been unclear. Our work provides the missing framework by positioning class A and class B within defined aggregation pathways and showing that their kinetic origins dictate both their toxic potency and whether they ultimately progress into fibrils. Together, these findings establish a mechanistic distinction: class B species follow the canonical fibril-surface generated oligomer route, whereas class A aggregates arise through an independent nucleation pathway. However, we cannot exclude that a small fraction of Class A species may enter fibril-forming pathways. We therefore interpret Class A as predominantly off-pathway and Class B as more closely linked to fibril-dependent secondary nucleation. Such on- and off-pathway intermediates have been reported for Aβ and other amyloidogenic proteins^68^, and species from both routes can contribute to toxicity and have been detected within Aβ plaques^69^. By contrast, class C species contribute minimally to overall membrane-disrupting activity only at the earliest stages of aggregation, and their formation route and molecular identity remain unclear.

After defining these aggregation routes *in vitro*, we next examined whether they also operate in the human brain to give rise to intermediates that contribute to neurodegeneration. Analysis of three AD brain extracts revealed diffusible, membrane-disruptive Aβ42 species corresponding to both class A and class B intermediates in every case examined, although their relative distribution varied between individuals. This variability may potentially reflect differences in disease stage, progression rate or genetic background. The coexistence of both intermediates across all cases supports a model in which primary and secondary nucleation pathways converge to drive pathogenic assembly in the human brain. Functionally, these species impaired human cortical neuronal connectivity and selective immunodepletion of Aβ42 significantly mitigated this deficit. Because this neurite outgrowth assay parallels deficits in hippocampal long-term potentiation^42^, a well-established correlate of learning and memory, these results implicate these Aβ42 intermediates as direct contributors to synaptic failure and cognitive decline in AD. Although the cohort size is limited, these data provide initial evidence that both intermediates are present in human AD brain, while larger studies will be required to define how their relative abundance varies across disease stages and patient populations. Together, these results establish a mechanistic bridge between aggregation routes defined in model systems and intermediates that persist in the human Alzheimer’s brain and highlight these intermediates as key drivers of neurodegeneration and promising targets for therapeutic intervention.

We put this therapeutic idea to the test in the *APP*^*NL-G-F*^ mice, which recapitulate key features of AD^22,51^. In BRICHOS R221E monomer-treated mice, plaque load was reduced, astrogliosis was attenuated, and cognitive function improved^52,53,70^, but the mechanistic basis for these improvements has remained unclear. BRICHOS does not block fibril formation outright; instead, it alters aggregation pathways, and how this rerouting translates into reduced pathology has not been resolved. To address this, we analyzed mouse brain homogenates and found that untreated (PBS-treated) mice contained both class A and class B intermediates, whereas BRICHOS-treated mice showed a marked reduction in the abundance of both populations. Class A intermediates were suppressed more strongly than class B, consistent with the differential sensitivity observed *in vitro*. These data indicate that BRICHOS acts by reshaping the oligomeric landscape *in vivo* and lowering the abundance of toxic intermediates.

These findings further show how eliminating intermediates attenuates multiple pathological axes simultaneously. At the neuronal level, Aβ42 intermediates exert direct toxicity by impairing connectivity; their suppression improve neurite integrity. In plaque formation, at least one class of intermediates serves as a precursor to fibrils and fuels plaque growth; reducing these intermediates decreases the pool of fibril-forming species and lowers plaque burden *in vivo*. Intermediates also induce inflammatory responses. Arctic Aβ42 intermediates triggered secretion of proinflammatory cytokines, providing a molecular explanation for the astrocytic activation observed in *APP*^*NL-G-F*^ mice. Exaggerated cytokine signalling causes synaptic and neuronal damage^71^ and sustains further reciprocal activation of microglia and astrocytes^72^. In addition, prolonged cytokine exposure impaired astrocytic uptake of Aβ, a major route of aggregate clearance^55^, paralleling earlier observations in microglia^73,74^ and explaining how intermediates can indirectly accelerate plaque accumulation. Finally, cytokine signaling also enhances neuronal Aβ production^75,76^, further amplifying aggregation burden and sustaining pathology. These data provide proof-of-principle that suppression of damaging Aβ intermediates *in vivo* correlates with disease-associated pathology.

Together, these processes converge into a self-reinforcing pathogenic cycle in which Aβ intermediates simultaneously drive neuronal injury, plaque accumulation, and pro-inflammatory signalling, each step further amplifying Aβ production and aggregation. Breaking this cycle requires suppressing intermediate formation at its point of origin, which reduces toxic assemblies, preserves neuronal health, limits inflammation, lowers plaque burden, and ultimately improves functional outcomes. These findings provide strong mechanistic evidence that early nucleation routes constitute a key actionable bottleneck through which the core pathology of AD can be directly intercepted.

Although these results advance our understanding of Aβ aggregation *in vivo*, several caveats remain. First, our assay detects species capable of membrane-disruption, so intermediates that do not perturb membranes may escape detection and some species may continue to evolve under the assay conditions. This selectivity, however, ensures that we capture only membrane-damaging aggregates, which are strongly implicated in AD^31–33^. Second, although our data show that these intermediates are functionally and biophysically distinct, higher-resolution structural analyses will be required to define how their molecular architectures give rise to their distinct activities. Third, post-translational modifications of Aβ42, particularly N-terminal truncation and pyroglutamate formation (Aβ_pE3_) are found in the AD brain^77^. In our study, recombinant Aβ_pE3_ produced two classes of intermediates with membrane-permeabilizing potency (Supplementary Fig. 15), similar to wild-type Aβ42. Because the C-terminus of Aβ42, a key determinant of membrane permeabilization^20^, remains unchanged in Aβ_pE3_, this likely accounts for their similar membrane-disrupting profiles to wild-type. The fact that Aβ_pE3_, like wild-type Aβ42, yielded two classes of species further supports the idea that nucleation-based aggregation routes operate in the human brain. Fourth, other Aβ isoforms are present *in vivo* and could complicate interpretation, most notably Aβ40, which is 10-20 times more abundant in healthy brain^78^. However, in regions most affected in AD, such as the cortex studied here, Aβ42 is predominant^18,19^, and we found Aβ42 levels were at least ten times higher than Aβ40 in both soluble aggregates (Supplementary Fig. 16) and insoluble plaques (Supplementary Fig. 8). Finally, while suppression of Aβ intermediates correlates with reduced plaque pathology and pro-inflammatory signalling in human cell models and Alzheimer’s mouse models, the contribution of these species to pathology extends beyond the mechanism examined here, and these findings should therefore be interpreted within the broader context of multiple Aβ-mediated pathogenic mechanisms^79^.

In summary, our findings demonstrate that toxic Aβ oligomeric intermediates are real, causal, and targetable in the Alzheimer’s brain. We define a mechanistic continuum between Aβ aggregation, plaque deposition, and neuronal toxicity across recombinant systems, mouse models, and the Alzheimer’s brain. By resolving aggregation pathways at the level of mechanism and function, our findings provide a framework for understanding how modulation of specific routes may limit the formation of toxic intermediates and contribute to reduced neuronal and glial pathology and plaque burden in model systems, supporting a testable strategy toward disease modification in AD. Detecting the same neurotoxic intermediates in Alzheimer’s brain as in our model systems supports the disease relevance of these models and highlights their value as platforms for therapeutic screening. More broadly, our framework establishes early molecular events as central drivers of disease progression and offers a generalizable strategy to intervene at the level of intermediate formation rather than downstream pathology, providing a path toward true disease-modifying therapies for early-stage AD and other protein-aggregation disorders.

## Methods

### Ethical statement

We complied with all relevant ethical regulations. All studies involving human and animal tissue were conducted in accordance with institutional guidelines and approved by the appropriate ethical review committees. Details of ethical approvals are provided in the corresponding Methods sections.

### Recombinant Aβ42 expression and purification

Recombinant Aβ42 was expressed from the pET-Sac-Aβ(M1-42) plasmid (Addgene, 71875) transformed into *E. coli* BL21 Star™ (DE3)pLysS One Shot™ Chemically Competent cells (ThermoFisher, C602003), as previously published^80^. Briefly, single colonies were selected on LB-agar supplemented with 100 µg/mL ampicillin (ThermoFisher, J60977.06) and expanded overnight in 1 L LB broth containing 100 µg/mL ampicillin. Then protein expression was induced with 1 mM IPTG (Neo Biotech, NB-45-00030-25g) at an optical density of OD600 = 0.45-0.5. Cells were incubated for 5h at 37 °C with shaking at 200 rpm before being pelleted by centrifugation at 6,000g for 10 min. Pellets were stored at -20 °C until further processing. For lysis, pellets were resuspended in buffer containing 10 mM Tris/HCl (pH 7.5) and 1 mM EDTA (Merck, 324503), supplemented with cOmplete™ EDTA-free protease inhibitor cocktail and DNase I (200 ng/mL) (Merck, 11873580001). Cells were lysed on ice by sonication (4 x 30s), followed by centrifugation at 18,000g for 8 min. This procedure was repeated twice. The resulting pellets were solubilized in 8M urea (Merck, U5378), 10 mM Tris/HCl (pH 8.5), 1 mM EDTA, and clarified by centrifugation at 18,000g for 5 min. The supernatant was diluted 1:4 in 10 mM Tris/HCl (pH 8.5) with 1 mM EDTA. Samples were applied to a gravity column (Cytiva, 19-5101-01) packed with 10 mL DEAE Sepharose beads (Cytiva, 17-0709-01), pre-equilibrated with 2M urea, 10 mM Tris/HCl (pH 8.5), 1 mM EDTA. The flow-through was collected and reapplied once, followed by three column washes with 10 mM Tris/HCl (pH 8.5) and 1 mM EDTA.. Aβ42 was eluted with 10 mM Tris/HCl (pH 8.5), 1 mM EDTA, 500 mM NaCl. Solid GuHCl (ThermoFisher, 24110) was added directly to a final concentration of 6 M and samples were filtered through a 0.2 µm membrane and incubated overnight at room temperature.

Size-exclusion chromatography (SEC) was performed using a HiLoad® 16/600 Superdex® 30 pg column (Cytiva, 28-9893-31) on an ÄKTA pure™ HPLC system, with 20 mM ammonium acetate and 0.2 mM EDTA (pH 8.0) as the running buffer. Fractions corresponding to the Aβ42 monomer peak were pooled, quantified using a Qubit™ Protein Broad Range Assay Kit (ThermoFisher, A50668), aliquoted, snap-frozen in liquid nitrogen, and stored at -80 °C.

### Arctic Aβ42 (E22G) Expression and purification

The Arctic Aβ42 (E22G) variant was generated from the plasmid pET-Sac-Aβ(M1-42) (Addgene, 71875) by site-directed mutagenesis using the QIAGEN QuickChange II kit according to the manufacturer’s protocol, with XL1-GOLD ultracompetent cells (Agilent, 200315) as hosts. Mutation-specific primers (forward 5′-ctggtgttcttcgctggagacgtgggttctaac-3′; reverse 5′-gttagaacccacgtctccagcgaagaacaccag-3′) were designed to introduce the E22G substitution. Mutated plasmids were purified using the QIAGEN Plasmid Miniprep kit, and DNA concentration and quality were verified by spectrophotometry. Then Arctic Aβ42 (E22G) was then expressed and purified similarly to wild-type Aβ42.

### Thioflavin T based aggregation assay

Aggregation kinetics of wild-type and Arctic Aβ42 were monitored using a Thioflavin T (ThT) fluorescence assay. Monomeric Aβ42 was diluted in phosphate-buffered saline (PBS) to 200 µM, aliquoted (50 µL), flash-frozen in liquid nitrogen, and stored at -80 °C. For aggregation assays, aliquots were thawed and further diluted in PBS to the desired concentration. Reactions were prepared in 96-well half-area plates (Corning, 3881) with 20 µM ThT (Sigma, T3516) and incubated at 37 °C without shaking. Plates were sealed, and fluorescence was monitored in real time using a Clariostar Plus plate reader (BMG Labtech).

For seeded aggregation assays, preformed Aβ42 fibril seeds were generated separately by incubating monomeric Aβ42 in PBS at 37 °C overnight. The resulting samples were transferred to low-binding tubes and sonicated for 2 min in a bath sonicator at room temperature. Fresh monomeric Aβ42 was then prepared as described above and supplemented with preformed fibril seeds at 1.5% or 2.5% relative to the monomer concentration. Seeded aggregation were performed in PBS containing 20 µM ThT at 37 °C under quiescent conditions.

N-terminally truncated and pyroglutamate-modified Aβ (Aβ_pE3_) was purchased (AnaSpec, AS-29907-01) and used without further purification. Aggregation assays were performed under identical conditions as for wild-type Aβ42. For BRICHOS R221E experiments, wild-type Aβ42 was aggregated in the presence of the chaperone at 10%, 50%, or 100% molar equivalents relative to monomer. For post-aggregation BRICHOS treatment experiments, pre-formed Aβ42 aggregates were collected at the aggregation midpoint and incubated with BRICHOS R221E at 100% molar equivalent relative to Aβ42 monomer before analysis using the single-vesicle Ca^2+^ influx assay. For small-molecule inhibition experiments, 3 μM Aβ42 was aggregated in the presence of N-(2-{[1,1′-biphenyl]-4-yl}ethyl)-3,4-dihydro-1H-2,3-benzoxazine-3-carboxamide (M1; Enamine, Z8478213987) at 0, 5, or 20 μM as previously mentioned^81^. Reactions were monitored with ThT fluorescence as described earlier.

### Dual-labelled Aβ42 aggregation

Monomeric Aβ42 conjugated to the FRET partner dyes HiLyte Fluor 555 (AnaSpec, AS-60480-01) and HiLyte Fluor 647 (AnaSpec, AS-64161) was obtained commercially. The peptides were first dissolved in 10 mM NaOH and subsequently purified by HPLC using a Bio-Sep 2000 column. Fractions corresponding to the peptide peak were collected, and concentrations were determined from the absorbance of the attached dye. Donor- and acceptor-labelled monomers were then combined in a 1:1 ratio so that half of the molecules carried the donor dye and half the acceptor dye. Aggregation reactions were initiated at a final protein concentration of 3 µM and incubated at 37 °C under quiescent conditions in PBS in 96-well half-area plates. Aliquots were taken at the same time points as those used for aggregation reactions performed with unlabelled Aβ42.

### Wide-Field Fluorescence Imaging Setup

Single-molecule and super-resolution imaging were performed using a custom-built microscope based on a Nikon Eclipse Ti2 body equipped with a Perfect Focus unit and Omicron Luxx lasers. Lasers were launched through a fibre coupler (KineFlex SM/PM Fiber), collimated with Zoom Fiber Collimators (Thorlabs C20FC-A), and passed through achromatic Quarter-Wave Plates (Thorlabs AQWP05M-600). Then the laser directed into the back focal plane of a 100x Plan ApoTIRF, 1.49 NA oil-immersion objective lens (Nikon). To achieve uniform illumination in epifluorescence and TIRF, a beam shaper (Asphericon, TSM25-10-LD-D-532) was integrated in the excitation path, resulting in less than 5% intensity variation across the imaging field^73^. Fluorescence emissions were collected through the same objective and separated using a dichroic beamsplitter. The light then passed through a specific set of optical filters for each fluorophore before being captured by a sCMOS camera (Photometrics Prime 95B). The whole setup was controlled by Micro-Manager 2.0.

### Vesicle preparation and immobilization in glass coverslips

Vesicles were prepared following established protocols with minor modifications^82^. Briefly, 35% PC (Avanti, 131101), 15% PE (Avanti, A85757) 8% PS (Avanti, A80534), 2% PI (Avanti, A85079) and 30% Cholesterol (Sigma, 700100P) and 10% Ganglioside GM1 (Avanti, A86065) were dissolved in chloroform and methanol mixture (4:1 v/v) and then further mixed with biotinylated lipids (18:1-12:0 Biotin PC; Avanti, 860563) at a molar ratio of 100:1. The solvent was removed under vacuum in a desiccator overnight, and the resulting lipid film was rehydrated in HEPES buffer (pH 6.5) containing 100 µM Cal-520 (500 µM for Arctic Aβ). The suspension underwent at least five freeze-thaw cycles, alternating between dry ice and a water bath. To obtain vesicles of ∼100 nm or smaller, the solution was extruded at least 10 times through a 100 nm pore-size polycarbonate membrane (Avanti). Unincorporated dye molecules were removed by size-exclusion chromatography using a Superdex™ 200 Increase 10/300 GL column on an ÄKTA Pure system (GE Life Sciences) with a flow rate of 0.5 mL/min.

For vesicle immobilization, borosilicate glass coverslips (22×22 mm^2^, VWR 6310122) were sequentially cleaned by sonication in Milli-Q water and methanol, dried under nitrogen, and plasma-treated for 20 min in an argon plasma cleaner (Deiner Zepto). Frame-Seal incubation chambers (9×9 mm^2^, Bio-Rad) were affixed to the coverslips, and surfaces were functionalised with a mixture of PLL-g-PEG and PLL-g-PEG-biotin (100:1, SuSoS AG; 1 g/L in HEPES buffer, pH 6.5) for 30 min. After washing with buffer, coverslips were incubated with NeutrAvidin (0.1 mg/mL in HEPES, ThermoFisher) for 15 min and washed again. Purified vesicles were then added to the chambers, incubated for 30 min, and excess vesicles were removed by three washes with reaction buffer.

### Membrane permeabilization assay

Membrane permeabilization was measured using a single-vesicle Ca^2+^-ion influx assay adapted from established protocols^50,83^. Vesicle chambers were first exchanged into Ca^2+^-containing buffer, and baseline fluorescence images (F_blank_) were acquired as 10 consecutive frames with 50 ms exposure per frame. These measurements provided the background intensity for each vesicle. Vesicles were then exposed to 1 nM Aβ42 (monomer equivalents), a concentration chosen to enable detection of discrete fluorescence increases corresponding to individual Ca^2+^-influx events within single vesicles within this time frame. After sample addition (F_sample_), imaging continued once every minute for 20 minutes using the same acquisition scheme: one imaging burst per minute consisting of 10 frames with 50 ms exposure each. Between imaging bursts, the sample remained unilluminated to minimise photobleaching. Following this period, ionomycin (1 mg/mL; Cambridge Bioscience, 1565) was added to fully saturate vesicles with Ca^2+^, and imaging proceeded for a further 5 minutes using the same settings (F_ionomycin_). Relative Ca^2+^-ion influx was calculated for each vesicle for each hit as:

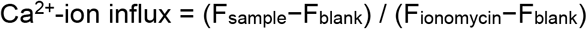

For analysis, the 10 frames within each imaging burst were averaged to obtain a single intensity value per burst. Under these conditions, most vesicles encounter Aβ aggregates and show discrete Ca^2+^ influx events without signal saturation. This averaging step improved the signal-to-noise ratio by reducing frame-to-frame fluctuations, allowing us to detect and quantify very small fluorescence changes corresponding to individual Ca^2+^-influx events. Cal-520 was excited at 488 nm, and emission was collected through a combination of a long-pass filter (BLP01-488R-25, Laser2000) and a band-pass filter (FF01-520/44-25, Laser2000) and imaged onto a Prime 95B sCMOS camera (Photometrics). Images were acquired at ∼1-10 W/cm^2^ power density with a scan speed of 20 Hz and 16-bit depth, using flat-field illumination with less than 5% intensity variation between the periphery and the centre of the image. Fields of view were selected automatically using a motorised stage controlled by Micro-Manager scripting to ensure unbiased data collection.

As controls, we incubated monomeric Aβ42 (1nM) with lipid vesicles in solution for 30 minutes and did not observe detectable aggregate formation (Supplementary Fig. 17). Similarly, when pre-formed aggregates (1nM) were incubated with lipid vesicles, the aggregate number remained unchanged. In addition, incubation of pre-formed Aβ42 aggregates (1 nM) in Ca^2+^ buffer alone showed no detectable change in aggregate number (Supplementary Fig. 18), indicating that neither the lipid membrane of the vesicles nor the Ca^2+^ buffer induces any artefacts under the experimental conditions.

### Vesicle size estimation using point accumulation for imaging in nanoscale topography (PAINT)

To determine the maximum fluorescence capacity of individual vesicles, we first estimated vesicle size in order to calculate the number of dye molecules encapsulated per vesicle, given that the dye concentration during vesicle preparation was known. Although a cutoff filter was used during preparation, it did not guarantee uniform size, as smaller vesicles could still form and introduce variation in dye content. For instance, a 100-nm cutoff filter can yield vesicles of both 100 nm and 50 nm in diameter; because volume scales with the cube of diameter, a 100-nm vesicle contains approximately eight times more dye molecules than a 50-nm vesicle.

Therefore, vesicle diameters were determined *in situ* using point accumulation for imaging in nanoscale topography (PAINT) with transient binding of Nile Red to vesicle membranes. First, Nile Red was dissolved in dimethyl sulfoxide and then diluted to 1 nM in PBS immediately prior to imaging. Samples were imaged in through-objective TIRF mode using a 532 nm laser for excitation. Emission was collected through a dichroic mirror (Di02-R532-25×36). Image sequences of 1,000 frames were acquired at 30 ms exposure. Localizations were extracted using the Peak Fit plugin for ImageJ/Fiji, and coordinates corresponding to individual vesicles were clustered using DBSCAN. Vesicle size was estimated by Gaussian fitting of the clustered localizations. Clusters containing fewer than 40 localizations were excluded as background. For calibration, fluorescence intensities obtained after ionomycin-induced Ca^2+^ influx were correlated with vesicle size.

### Analogue Ca^2+^-ion influx conversion to digital number of Ca^2+^-ion influx

To convert the analogue percentage readout of Ca^2+^-ion influx into absolute ion numbers, we calculated the number of dye molecules per vesicle from vesicle size data. The number of Cal-520 per vesicle was estimated from the known encapsulated Cal-520 concentration and the vesicle volume, derived from PAINT-based diameter measurements. The theoretical dye number was then used to convert percentage influx values into ion counts according to:

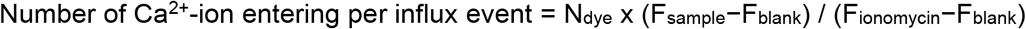

where N_dye_ is the number of Cal-520 molecules per vesicle. The fractional values obtained were rounded to yield discrete Ca^2+^ ion counts. From these measurements, we calculated the number of dye molecules encapsulated in each vesicle. Experiments were conducted within a range where Cal-520 fluorescence intensity varied linearly with concentration, avoiding dye aggregation or quenching. We measured the encapsulation efficiency (EE), defined as the ratio between measured and calculated intensity, by plotting integrated Cal-520 fluorescence against vesicle volume. In our system, EE was close to unity for vesicles up to ∼100 nm in diameter but decreased for larger vesicles. Therefore, vesicles were prepared with a 100 nm cutoff filter, and vesicles larger than this threshold were excluded from analysis. Effective vesicle diameters were then obtained by subtracting 10 nm from the measured diameters to account for bilayer thickness^84^. Similar to the previous report, we also found empty vesicles that did not contain any encapsulated dye^37^, and these were excluded from the analysis.

For control experiments, BRICHOS R221E alone was used at the same concentrations and incubated under the same conditions as in the Aβ42 aggregation reactions. The samples were incubated in the same well plate for the same duration before being added to vesicles immobilized on the coverslip. Membrane permeabilization measurements were then performed using the same assay conditions and imaging procedure as described for the Aβ42 samples.

### Expression and Purification of Bri2 BRICHOS R221E Monomer

The Bri2 BRICHOS R221E variant was prepared following a previously described protocol with minor modifications^40,64^. Site-directed mutagenesis was performed on the wild-type His_6_-NT*-Bri2 BRICHOS (residues 113-231) plasmid to introduce the R221E substitution. The construct was expressed in *E. coli* Shuffle T7 cells cultured in LB medium supplemented with 15 µg/mL kanamycin at 30 °C. Protein expression was induced with 0.5 mM IPTG at an optical density of 0.8, and cultures were incubated overnight at 20 °C. Cells were harvested by centrifugation (5,000g, 20 min, 4 °C), resuspended in 20 mM Tris-HCl buffer (pH 8.0), lysed by sonication, and clarified by centrifugation (24,000g, 30 min, 4 °C). The NT*-BRICHOS fusion protein was purified by Ni-NTA affinity chromatography. To remove the His_6_-NT* solubility tag, thrombin digestion was performed during overnight dialysis, leaving an additional glycine and serine at the N-terminus of BRICHOS. A second Ni-NTA purification step was used to separate the His_6_-NT* fragment, and the flow-through containing BRICHOS was collected. Samples were heat treated at 60 °C for 10 min and monomeric protein was isolated by size-exclusion chromatography on a HiLoad Superdex 75 pg column. The purified BRICHOS R221E monomer was concentrated to the desired level and stored at -70 °C until use.

### Endotoxin measurement in recombinant protein

Endotoxin levels in recombinant Aβ42 and Bri2 BRICHOS R221E monomer preparations were quantified using a commercial endotoxin detection kit (Thermo Fisher Scientific, Cat. No. A39553) according to the manufacturer’s instructions. Endotoxin levels were 0.03 EU/mg for recombinant Aβ42 and 0.05 EU/mg for Bri2 BRICHOS R221E monomer. Furthermore, recombinant Aβ42 was passed through an endotoxin removal spin column (ThermoFisher, Cat. No. 88274). The protein concentration was remeasured and aggregation were performed using ThT, which remained unchanged (**Figure S19A)**. To further assess potential functional consequences of endotoxin contamination, we analyzed astrocyte-mediated cytokine secretion using the resulting aggregates, as astrocytes are highly sensitive to even minimal levels of endotoxin. Following endotoxin removal, we observed no differences in astrocyte cytokine responses compared with untreated samples (**Figure S19B**).

### Basis function fitting

Basis functions representing the Ca^2+^ ion influx distribution of class A, B and C intermediates were selected as described in the Results. These were then normalized to unity and fitted using a simple python script to the overall Ca^2+^ ion influx distributions at each time point and condition using a standard nonlinear least squares method.

### Kinetic modeling of fibril formation

The kinetics of fibril formation by Aß42 can be well-described by a model featuring three reaction steps. These are primary nucleation of new fibrils with rate constant *k*_*n*_ and reaction order with respect to monomer *n*_*c*_, fibril surface-catalyzed secondary nucleation with rate constant *k*_2_ and reaction order with respect to monomer *n*_2_, and elongation with rate constant *k*_+_. The integrated rate law corresponding to this model was determined in^85^. In this model, fibril mass concentration (which is the quantity tracked by ThT fluorescence) is denoted *M*(*t*), and total initial monomer concentration is denoted *m*_tot_. Setting the saturation concentrations to infinity, which are not needed here due to the relatively small concentration range of 2-5 µM, gives:

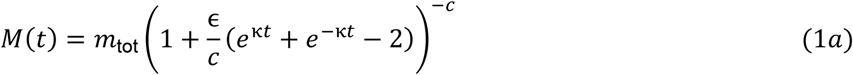

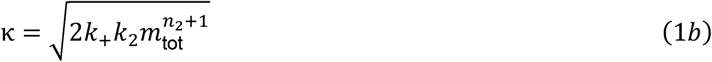

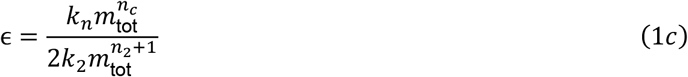

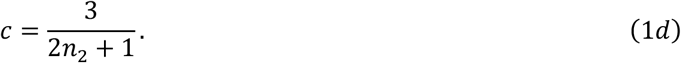

Fitting the data with this model gives combined rate constants 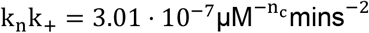, 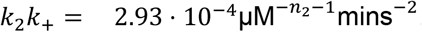, and reaction orders *n*_*c*_ = 1.74, *n*_2_ = 1.37. The *n*_2_ value <2 reflects partial saturation; Aß42 secondary nucleation is known to saturate as *m*_tot_ approaches 6 µM. Fits were performed using an offline version of AmyloFit^86^, which is largely python-based.

### Kinetic modeling of oligomers

The basis function fitting produced kinetic curves for class A and B intermediates at 3 distinct initial monomer concentrations. Oligomer kinetics can be described with the following integrated rate laws^46^, where *S*(*t*) denotes oligomer concentration:

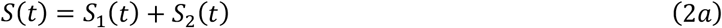

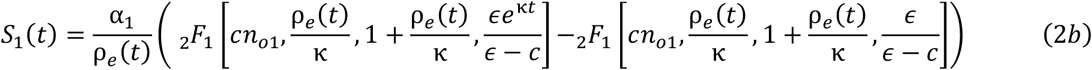

where _2_F_1_[a,b;c;z] is the Gaussian hypergeometric function.

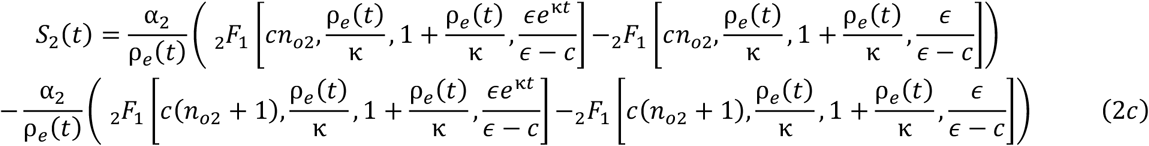

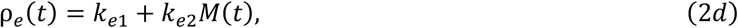

With M(t) given by Equation (1). The parameters α_1_, α_2_ are given by:

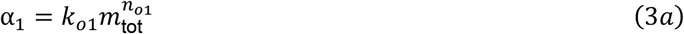

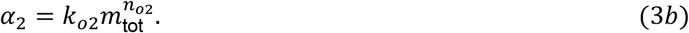

Since dissociation must be catalyzed by fibril surfaces comparably to oligomer formation, the additional relation holds:

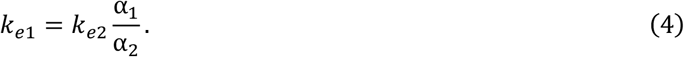

If the oligomers are on-pathway to fibril formation, the rate constants from the 2-state model used to fit the fibril kinetic curves can be used to constrain the rate constants in the oligomer kinetic model using the following formulae from^46^:

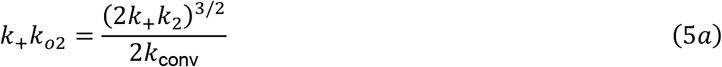

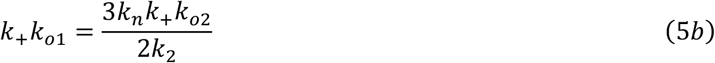

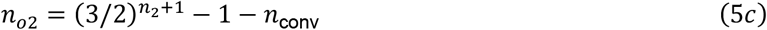

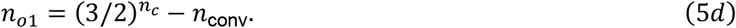

Using these relations, the rate equations can then be fitted to the oligomer kinetic curves, fitting the free parameters *k*_conv_, *n*_conv_, *k*_*e*2_. The remaining parameters *k*_+_*k*_*o*1_, *k*_+_*k*_*o*2_, *n*_*o*1_, *n*_*o*2_ are then calculated using Eqs. (5) after fitting.

If an oligomer population is instead off-pathway, equation (5) do not apply and the free parameters for fitting are instead *k*_+_*k*_*o*1_, *k*_+_*k*_*o*2_, *n*_*o*1_, *n*_*o*2_, *k*_*e*2_.

Again, fitting was performed using an offline implementation of AmyloFit^86^. (Note, since the number of vesicle binding events per nM of oligomers is unknown, the fitted *k*_*o*1_, *k*_*o*2_ are valid only up to an unknown constant of proportionality.) This yielded misfits in the case that class A intermediates are considered on-pathway. Good fits were instead obtained when treating class A intermediates as off-pathway, with the following combined rate parameters: *k*_*e*2_ = 0.072 mins^−1^, *n*_*o*1_ = 2.19, *n*_*o*2_ = 0.606, and 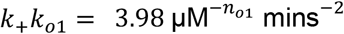, 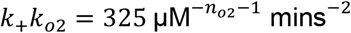.

By contrast, good fits were determined when class B intermediates were considered on-pathway, with the following rate parameters: *k*_*e*2_ = 0.00101 mins^−1^, 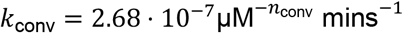, *n*_conv_ = 0.44.

### Human postmortem tissue case selection

Postmortem human brain tissue was obtained from the Sheffield Brain Tissue Bank (SBTB), University of Sheffield, which operates under ethical approval as a Research Tissue Bank from the Wales Research Ethics Committee 5 (Reference 24/WA/0052). Informed consent was obtained from all donors during life or from their next of kin. Tissue samples were collected from three individuals (Female, 92 years; Female, 79 years; Male, 71 years) with clinically diagnosed and pathologically confirmed AD. To assess both amyloid plaque pathology and the presence of soluble Aβ assemblies, we focused on cortical regions that consistently display substantial Aβ deposition in AD. From each donor, two complementary tissue preparations were obtained from the *same anatomical areas*: *(i)* formalin-fixed, paraffin-embedded (FFPE) samples for immunohistochemical analysis, and *(ii)* flash-frozen tissue for biochemical extraction of diffusible aggregates. FFPE tissue is required for plaque quantification, whereas flash frozen tissue preserves the native conformations of soluble Aβ species, which are altered by formalin cross-linking. To allow these parallel analyses, each brain was divided sagittally. One hemibrain underwent formalin fixation before dissection and paraffin embedding, while the opposite hemibrain was sectioned coronally and immediately flash-frozen in liquid nitrogen-cooled teflon-coated copper plates.

### Immunohistochemistry of postmortem AD tissue

Formalin-fixed paraffin-embedded (FFPE) frontal cortex sections (5 µm) from three AD cases were used for immunohistochemistry with an antibody specific for Aβ40 (EPR23712-2, Abcam), Aβ42 (21F12, Absolute Antibody, Ab02391-23.0) and DAPI. Sections were dewaxed in xylene, rehydrated through a graded ethanol series, and incubated in 3% H_2_O_2_ in methanol for 10 min to quench endogenous peroxidase activity. Antigen retrieval was performed in citrate buffer (pH 6.0) by microwave at full power for 10 min, followed by a 3 min incubation in 100% formic acid. To block nonspecific binding, sections were incubated with 2.5% normal horse serum (VECTASTAIN^®^ Elite ABC-HRP Kit, Mouse IgG, Vector Laboratories, PK-6102) for 30 min at room temperature. Sections were then incubated overnight at 4 °C with primary antibody (1:500 and 1:1000 dilution respectively), followed by Alexa Fluor 488 (Thermo Fischer, A-11008) and Alexa Fluor 647 (Thermo Fischer, A-11004) secondary antibody for 1 hour at room temperature. Slides were scanned at 20x magnification using a Hamamatsu NanoZoomer XR slide scanner.

### Extraction of diffusible protein aggregates from human postmortem tissue

Diffusible Aβ-containing aggregates were extracted from postmortem human frontal cortex tissue using a previously described protocol with minor modifications^48,49^. Briefly, tissue samples were cut into ∼300 mg pieces with a razor blade and incubated with 1 mL artificial cerebrospinal fluid (aCSF; 120 mM NaCl, 2.5 mM KCl, 1.5 mM NaH2PO4, 26 mM NaHCO3, 1.3 mM MgCl2, pH 7.4) at 4 °C for 30 min under gentle agitation. Samples were centrifuged at 2,000g for 10 min at 4 °C, and the upper ∼80% of the supernatant was collected and centrifuged again at 14,000g for 2 h at 4 °C. The resulting supernatant (∼80% of volume) was dialyzed for 72 h at 4 °C against a 50-fold excess of aCSF using a 2 kDa molecular weight cutoff (MWCO) Slide-A-Lyzer (Thermo Fisher), with buffer exchanges every 12 h. Dialyzed extracts were aliquoted, snap-frozen, and stored at -80 °C until use, avoiding additional freeze-thaw cycles.

### Immunohistochemistry of *APP*^*NL-G-F*^ mouse tissue

Brain tissue from *APP*^*NL-G-F*^ knock-in mice that had been treated previously^52,53^, with either PBS or recombinant human BRICHOS R221E was used for analysis. In that study, animals received intravenous injections of PBS (vehicle) or BRICHOS R221E (10 mg/kg) twice weekly for 12 weeks beginning at 9 months of age, followed by a four-week treatment-free period. At the study endpoint, brains were dissected and processed for both immunohistochemistry and biochemical assays. For histological analyses, the hippocampus and retrosplenial cortex from the right hemispheres were immersion-fixed in 4% paraformaldehyde (PFA) in 0.1 M phosphate buffer at 4 °C overnight, cryoprotected in 30% sucrose for 24 h, and transferred to cryoprotectant solution at -20 °C. Coronal sections (20 µm) were cut and used for immunostaining. Sections were washed in PBS, permeabilized with 0.3% Triton X-100 in PBS for 20 min at room temperature, and blocked for 30 min in PBS containing 0.1% Triton X-100 and 5% normal donkey or goat serum. Primary antibodies included anti-GFAP (rabbit, 1:800; Dako, Z033401-2) and anti-Aβ (mouse monoclonal 82E1, 1:200; IBL, 10323). Sections were incubated overnight at 4 °C, washed in PBS, and then incubated for 1 h at room temperature with Alexa Fluor Plus 594 donkey anti-mouse IgG (1:400; Invitrogen, A32744) or swine anti-rabbit IgG (1:200; Dako, P0399). Sections were washed again, mounted with ProLong Glass Antifade Mount (Invitrogen, P36984), and imaged using a Nikon Eclipse E800 microscope. All animal handling and procedures followed the comparative medicine guidelines of Karolinska Institutet, under ethical approval from the Stockholm Ethical Committee for Animal Experiments.

### Preparation of *APP*^*NL-G-F*^ mouse brain homogenates

After the brain dissection, the hippocampus and cerebral cortex of the left hemispheres were frozen on dry ice and stored at -80 °C. These fresh-frozen tissue was homogenized in Tris buffer (20 mM Tris-HCl, 150 mM NaCl, pH 7.4) supplemented with protease and phosphatase inhibitors (Thermo Fisher, 78440). Homogenates were centrifuged at 50,000 rpm for 20 min at 4°C, and the resulting supernatant was collected as the TBS fraction. The pellet was resuspended in Tris buffer containing 1% Triton X-100, homogenized, and centrifuged under the same conditions to generate the TBST fraction. For each treatment group (PBS and BRICHOS), homogenates from six mice were pooled within each treatment group, total protein concentrations were determined, and aliquots were stored at -80 °C until use.

### Immunoprecipitation (IP) assay

IP was performed using the Dynabeads Protein G Immunoprecipitation Kit (Thermo Fisher, 80105G). To minimize nonspecific binding, 5 µL of Protein G beads were washed three times with the kit-supplied buffer and incubated with human or mouse brain extracts for 30 minutes at room temperature. This pre-clearing step was performed without antibody, and the resulting supernatant was collected for immunodepletion. For antibody coupling, 50 µL of Protein G coated Dynabeads were incubated with 5 µg of Aβ42-specific 21F12 antibody or isotype control (Thermo Fisher, 02-6300) for 1 hour at room temperature. Beads were then placed on a magnetic rack, and the supernatant was discarded. After a single wash with 200 µL of wash buffer, antibody-bound beads were mixed with the pre-cleared brain extracts and rotated overnight at 4 °C to allow immunoprecipitation. The next day, samples were centrifuged at 6,000 rpm for 3 minutes at 4 °C. Approximately 80% of the supernatant was collected, aliquoted, and stored at -80 °C in low-binding protein tubes for downstream analysis.

### Antibody labelling for SiMPull assay

The 21F12 antibody was conjugated with Biotin or Alexa Fluor 647 using a Lightning-Link® conjugation kit (Abcam) according to the manufacturer’s instructions. In brief, 10µL of antibody was mixed with 1µL of modifier reagent and transferred to the lyophilized conjugation mix. The reaction was incubated for 15 min at room temperature in the dark, followed by addition of 1µL quencher reagent. After a further 5 min incubation, the labelled antibody was stored at 4°C until use.

### Single-molecule pull-down (SiMPull) assays for detecting Aβ42 aggregates in brain extracts

SiMPull assay was performed on covalently PEGylated glass coverslips (VWR, MENZBC026076AC40) as previously described^49^, with minor modifications. Assay wells were first coated with 10 µL of NeutrAvidin (Thermo Fisher, 31000; 0.2 mg/mL in PBS containing 0.05% Tween-20) for 5 minutes, washed, and incubated with biotinylated 21F12 antibody (10 nM in PBS) for 15 minutes. After washing, recombinant protein preparations or human/mouse brain extracts were added and incubated for 1 hour at room temperature. Bound Aβ species were detected using either Alexa Fluor 647–labeled 21F12 antibody (1 nM in PBS) or anti-pyroglutamate antibody (Abcam, IBR-10-2-3), incubated for 30 minutes. Between each step, wells were washed with PBS containing 0.05% and 1% Tween-20. Finally, 3 µL of PBS was added to each well, and samples were sealed with a second plasma-cleaned coverslip.

### Super-resolution imaging of Aβ aggregates

Super-resolution imaging of recombinant Aβ42 aggregates was performed using direct stochastic optical reconstruction microscopy (dSTORM) combined with the SiMPull approach. Aβ42 aggregates were first captured on the surface using biotinylated 6E10 antibodies and subsequently detected with Alexa Fluor 647–labelled 6E10 antibodies. Following the imaging antibody incubation step of the SiMPull protocol, the PBS buffer was removed and replaced with a freshly prepared dSTORM imaging buffer containing 1 mg/mL glucose oxidase, 50 µg/mL catalase, and 50 mM cysteamine in 50 mM Tris buffer supplemented with 10% (v/v) glucose in PBS (pH 8), which served as an oxygen-scavenging system. Images were acquired using 647 nm laser excitation with an exposure time of 30 ms, and a total of 3,000 frames were recorded for each acquisition. Localization of individual blinking events was performed using the Peak Fit module of the GDSC Single Molecule Light Microscopy plugin for ImageJ. Localizations were filtered using a signal intensity threshold of 40 a.u. and a localization precision threshold of 30 nm. Image reconstruction and analysis were carried out as described previously^87^.

### Separation of Aβ42 aggregates by sucrose gradient and ThT-based single-aggregate imaging

Aliquots of 3 µM Aβ42 aggregation reactions were collected at 85 minutes, when both class A and class B species are present. For fractionation, 400 µL of the aggregation mixture was layered onto a sucrose step gradient ranging from 10% to 50% (w/v) in PBS, prepared in 10% increments. The density of the sucrose solutions was verified by refractive index measurements. Gradients were centrifuged for approximately 5h at ∼25,000g at room temperature. Fractions were collected immediately after centrifugation and used directly for subsequent analyses.

For ThT-based single-aggregate imaging^23^, fractions corresponding to 10-20% and 30-40% sucrose were mixed separately. Glass coverslips were first plasma-cleaned using an argon plasma cleaner for 20 min and subsequently coated with poly-L-lysine (Merck, P4707) for 20 min. The collected fractions were mixed with 5 µM ThT and added to the coverslips. After a 15 min incubation, imaging was performed using TIRF microscopy.

Surface-exposed hydrophobicity of Aβ aggregates was assessed using 8-anilino-1-naphthalenesulfonic acid (ANS) fluorescence. Aβ aggregate samples were diluted to a final concentration of 4 μM monomer equivalent and incubated with 40 μM ANS in 96-well half-area plates. Fluorescence spectra were recorded using a Clariostar plate reader with excitation at 350 nm and emission collected from 420-600 nm.

### Influence of Ca^2+^ buffer and lipid vesicles on Aβ42 aggregation under assay conditions

To assess whether the Ca^2+^ imaging buffer used in the membrane permeabilization assay alters the aggregate population, pre-formed Aβ42 aggregates (3 µM aggregation, collected at 85 min) were transferred into the Ca^2+^ buffer and incubated for up to 30 min. Aggregate number was quantified at the beginning and end of the incubation using single-aggregate imaging.

To evaluate potential effects of lipid membranes, monomeric Aβ42 or pre-formed aggregates (3 µM aggregation, collected at 85 min) were incubated separately with lipid vesicles in Eppendorf tubes using the same vesicle composition and concentrations as employed in the membrane permeabilization assay. After incubation, vesicles were lysed with 0.5% Tween-20, and the samples were analyzed using the same single-aggregate detection method to quantify aggregate abundance.

### Differentiation of NPC into cortical neurons and its characterization

Neural progenitor cells (NPCs) and cortical neurons were derived from iPSCs following a published protocol with minor modifications^88^. The iPSC line KOLF2.1J (JIPSC001162) was obtained from The Jackson Laboratory and used in accordance with the supplier’s usage terms.. This is an control line carrying the *APOE3/3* genotype, which is considered neutral with respect to AD risk. Since our aim was to assess the neurotoxic potential of Aβ aggregates, we selected this *APOE3/3* control line to generate healthy neurons that provide a neutral background. Cells were thawed, maintained in StemFlex medium (Gibco, A3349401) on Matrigel-coated plates, and passaged using ReLeSR (STEMCELL Technologies, 100-0484). Colonies were regularly examined to confirm typical iPSC morphology, and pluripotency was verified by immunostaining for established markers. Neural induction was initiated when cultures reached ∼70-80% confluency. iPSCs were dissociated to single cells with Accutase, replated at high density, and cultured in neural induction medium containing dual-SMAD inhibitors SB431542 (STEMCELL Technologies, 72234) and LDN-193189 (Selleckchem, DM3189), alongside Rock inhibitor to promote survival. Within the first week, cells formed a confluent neuroepithelial layer, and neural rosettes typically emerged between DIV12-15. Rosettes were manually selected and expanded using the STEMdiff Neuronal Rosette Selection Reagent (STEMCELL Technologies, 05832). Cultures were maintained in neural maintenance medium, with or without bFGF supplementation depending on the stage of rosette development. Rosettes were routinely frozen between DIV35-41 to establish working stocks, and thawed as needed for further differentiation.

For cortical neuron generation, neural rosettes were dissociated at DPT7 and replated onto poly-L-ornithine/laminin-coated plates in neuronal plating medium supplemented with DAPT and RevitaCell. Neurons were then maintained in Neurobasal Plus medium with B27, GlutaMAX, and antibiotics. Proliferating cells were suppressed by the addition of 5-fluorouracil/uridine (5FU) starting from DIV3, whilst DAPT was maintained for 7 days post-plating. From DIV7, the medium was supplemented with the neurotrophic factors BDNF (Qkine, Qk050-1000) and GDNF (Qkine, Qk051-1000) (10 ng/mL each) to support maturation and survival. Cells were maintained with half-medium changes two to three times per week. Mature neuronal networks were established by DIV28-30.

Cells cultured in 96-well plates were washed with PBS and fixed in 3.7% paraformaldehyde for 20 min at room temperature. Following permeabilization with 0.2% Triton X-100/PBS for 10 min, nonspecific binding was blocked using 0.2% fish gelatine (Merck, G7765) in PBS for 1 h. Primary antibodies were diluted in blocking solution and incubated with cells overnight at 4°C: Nanog (Cell Signaling, 4903; 1:500), Pax6 (BioLegend, 901302; 1:500), FoxG1 (Invitrogen, PA5-2679; 1:25), Ctip2 (Abcam, ab18465; 1:300), βIII-tubulin (Abcam, ab18207; 1:1000), and MAP2 (Abcam, ab5392; 1:5000). The following day, cells were washed and incubated for 1h at room temperature with Alexa Fluor-conjugated secondary antibodies (Invitrogen; 1:500). Nuclei were counterstained with Hoechst 33342 (Life Technologies, H3570, 1:5000;) for 10 min. After final washes in PBS, stained cells were imaged using an Opera Phenix™ High Content Imaging System (PerkinElmer).

### Live-neuronal imaging for neurite length measurement

To assess the neurotoxic effects of Aβ aggregates, hPSC neurons were plated in 96 well plates at a density of 30,000 cells/ per well and placed in an IncuCyte live-cell imaging system and monitored for 5 days. The Aβ42 aggregates used for this experiment were collected from the same aggregation reactions used for the membrane permeabilization assays. On DIV28 and DIV30, neurons were treated with 0.1µM Aβ42 oligomers (either alone or prepared in the presence of BRICHOS R221E) at 48 h intervals. Cultures were imaged every 6 hours, 9 different area are per well. Neuronal network integrity was quantified as neurite length using the IncuCyte® NeuroTrack Analysis Module. A schematic of the neuronal culture timeline, quantification of key markers and treatment paradigm for neurite outgrowth assay is shown in Supplementary Fig. 20.

### iAstrocyte culture and response to Arctic Aβ42

iAstrocytes were generated from fibroblasts obtained from a healthy control (female, age 72) via an induced neuronal progenitor cell (iNPC) intermediate, following previously published differentiation protocols^89^. This line was obtained through the MODEL-AD research study (Research Ethics Committee number: 16/YH/0155). Fibroblasts were plated at 250,000 cells per well and transduced for 12h with retroviral vectors encoding OCT3, SOX2, and KLF4. After 48 h, cultures were transitioned into iNPC medium consisting of DMEM/F12 (Thermo Fisher, 11320033), 1% N2 supplement (Thermo Fisher, 17502048), 1% B27 (Thermo Fisher, A3653401), and FGF (40 ng/mL, Peprotech, 100-18B). To promote astrocytic differentiation, iNPCs were seeded onto fibronectin-coated (5 µg/mL; R&D Systems, 1918-FN) 10 cm dishes in DMEM media with high glucose (ThermoFisher, 11965092) supplemented with 10% FBS (Thermo Fisher, 16000044), 0.2% N2 for 7 days and 50 U/mL of Penicillin-Streptomycin (ThermoFisher, 15140122). iAstrocytes were candidates using established markers vimentin, CD44, and EAAT2. Primary antibodies included anti-vimentin (Merck Millipore, AB5733; 1:1000), anti-CD44 (Abcam, ab157107; 1:200), anti-EAAT2 (Abcam, ab41621; 1:250), and GFAP (Agilent Dako, Z0334, 1:200) diluted in PBS with 1% BSA and 0.1 mg/mL horse serum. Secondary antibodies were Alexa Fluor 488 goat anti-chicken IgG (ThermoFisher, A-11039; 1:1000) for vimentin, and Alexa Fluor 488 donkey anti-rabbit IgG (ThermoFisher, A-21206; 1:1000) for CD44, EAAT2 and GFAP. A schematic of the iAstrocyte culture timeline, marker quantification, and treatment paradigm for the Aβ uptake assay and cytokine measurements in conditioned media is shown in Supplementary Fig. 21.

For experiments with Arctic Aβ42, iAstrocytes were plated at 2,500 cells per well in 96-well plates (Greiner Bio-One) and exposed to Arctic Aβ42 aggregates (1 µM monomer equivalents). Cells were incubated for 1h, after which medium was collected, snap-frozen in liquid nitrogen, and stored at -80°C for cytokine analysis. Following exposure, cells were fixed with 4% formaldehyde for 10 min and permeabilized with 0.1% Triton X-100 (Sigma-Aldrich, X100-100ML) in PBS-T (0.1% Tween-20 in PBS) for 30 min, followed by two washes in PBS-T. To block nonspecific binding, cells were incubated in PBS-T containing 5% horse serum for 1h. Primary antibodies against CD44 (Abcam, ab157107) and Aβ (6E10; BioLegend, 803001) were applied in PBS with 1% BSA and incubated overnight at 4°C. Secondary antibodies Alexa Fluor 488-labelled goat anti-mouse IgG (Thermo Fisher, A-11029) and Alexa Fluor 647-labelled donkey anti-rabbit IgG (ThermoFisher, A-31573) were added at 1:1000 for 1 h, followed by nuclear counterstaining with 2.5 µM Hoechst (Life Technologies, H3570) for 10 min. iAstrocytes were washed three times in PBS, and internalized Aβ42 was imaged using the Opera Phenix High Content Imaging System (PerkinElmer).

### ELISA to measure cytokine in iAstrocytes media

To assay cytokine secretion by iAstrocytes, cell media were collected after incubation of cells with Aβ samples for 1 h and stored at -80 °C until analyzed. IL-1β, IL-6 and TNF-α in the cell media were measured using the Duoset® enzyme-linked immunosorbent assay (R&D Systems DY201, DY206, DY210), according to the manufacturer’s protocol.

### Statistical tests

No statistical method was used to pre-determine sample size and no data were excluded from the analyses. All membrane permeabilization assays were performed in three independent biological replicates, which were pooled to obtain sufficient event numbers for reliable kinetic fitting. Cellular experiments were carried out in triplicate biological replicates. Statistical significance between two groups was determined using unpaired two-tailed t-tests, whilst comparisons across multiple groups were made using one-way ANOVA followed by Tukey’s post hoc test. Comparisons with *P* < 0.05 were considered statistically significant and are indicated with asterisks (\**P* < 0.05, \*\**P* < 0.01, \*\*\**P* < 0.001 and \*\*\*\**P* < 0.0001). All statistical analyses were performed using Origin 9.0 (OriginLab).

## Data availability

All data are available within the article and supplementary information and a are available from the corresponding authors upon request.. Source data are provided with this paper.

## Code availability

No new code or software was created for this manuscript. Procedures for mathematical modeling and statistical analysis of data are described in Methods.

## Acknowledgments

We gratefully acknowledge the Sheffield Brain Tissue Bank for providing tissue samples, and we extend our sincere thanks to the donors and their families for their invaluable contribution to scientific research. We are grateful to Tuomas P.J. Knowles, University of Cambridge for a stimulating discussion regarding our manuscript.

## Author contributions

A.U., D.P.R., H.E.W., E.F.G., and H.J.B. performed the majority of the experiments. A.D. and G.M. designed and conducted the kinetic modeling. A.U. and E.F.G. carried out the human hiPSC neuron studies, whilst H.E.W. and S.B. performed the iAstrocyte experiments. D.P.R. and H.J.B. conducted the *in vitro* protein assays. J.R.H. selected and provided human postmortem tissue samples. Histopathological analysis and tissue homogenization were performed by E.F.G. and A.U. W.H.M. expressed and purified BRICHOS R221E, and R.Z. conducted the immunohistochemistry of mouse tissue. A.A., G.C., S.M.B., J.A.V., T.C.T.M., J.R.H., G.M., J.J., and S.D. supervised the work. The initial draft of the manuscript was prepared by A.U., A.D., G.M. and S.D., with all authors reviewing, providing critical feedback, and approving the final version. The study was conceived and designed by S.D.

