## Supporting info file for "Two functionally distinct amyloid-β intermediates link aggregation pathways to pathology in Alzheimer’s disease"

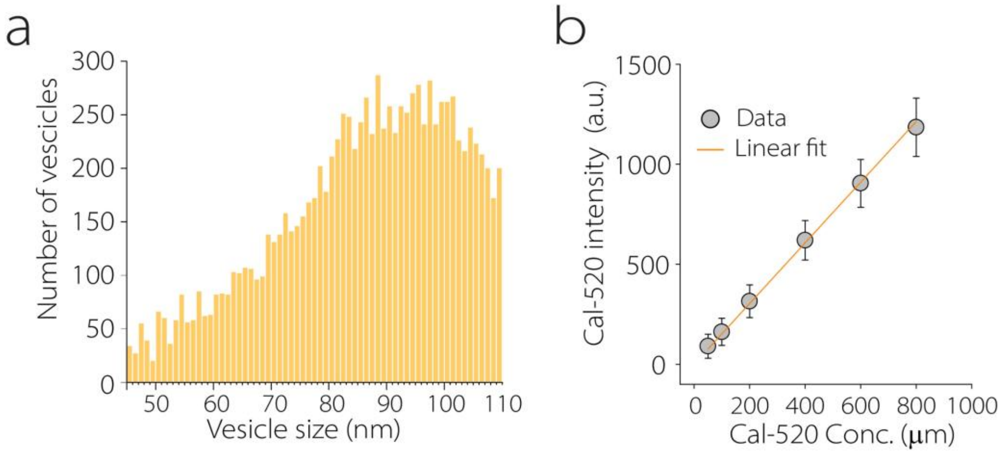


**Supplementary Figure 1. (a)** Size distribution of individual lipid vesicles measured using Nile Red PAINT **(b)** Cal-520 fluorescence intensity as a function of dye concentration. Data points (grey circles) represent mean intensity values with standard deviation, and the linear fit (orange line).

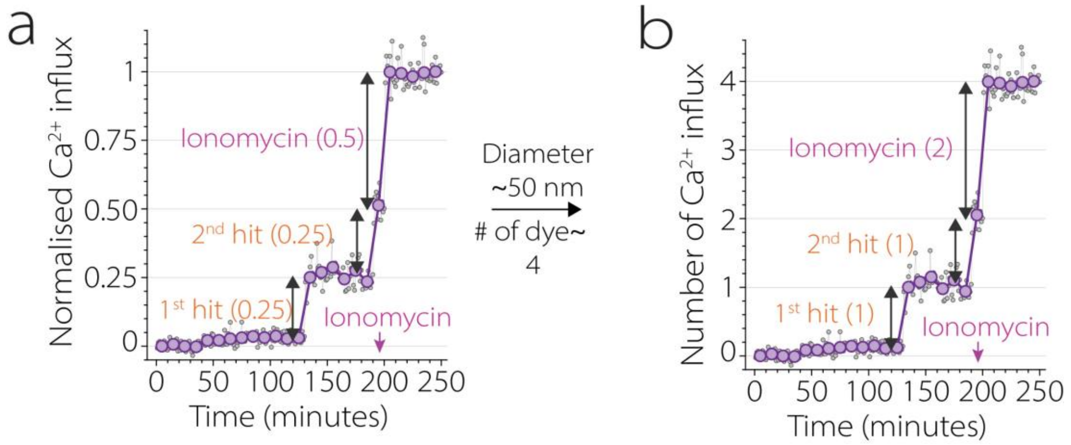


**Supplementary Figure 2. (a,b)** Fluorescence traces of $Ca^{2+}$ influx measured from a single vesicle (~50 nm diameter) shown in analogue **(a)** and digital **(b)** representations. Small grey circles indicate the fluorescence intensity of individual frames within each 10-frame imaging burst, while purple circles represent the averaged intensity of each burst acquired once per minute. A "hit" refers to a detected $Ca^{2+}$ influx event.

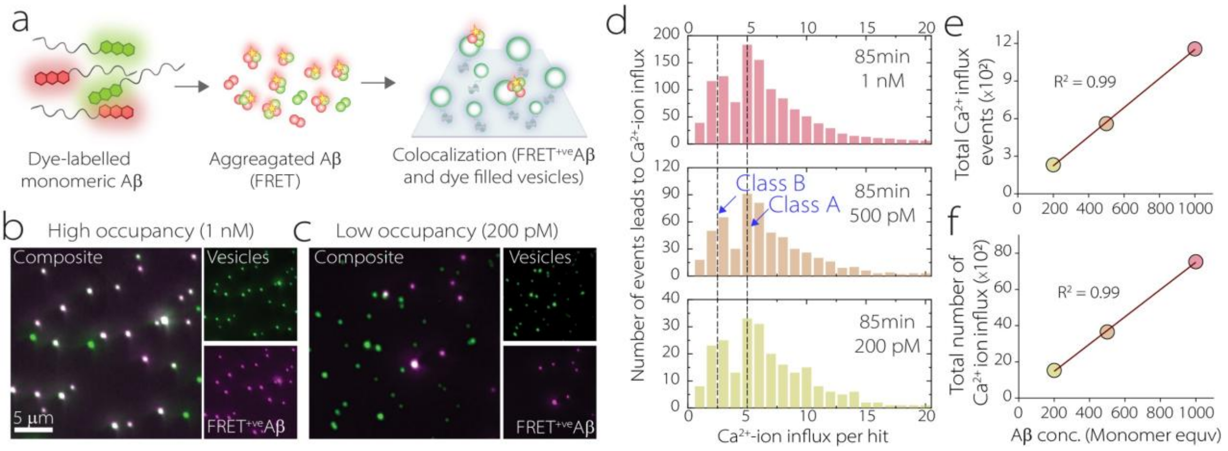


**Supplementary Figure 3.** (**a**) Monomeric Aβ labeled with HiLyte Fluor 555 and HiLyte Fluor 647 were mixed 1:1 and aggregation were performed. Aggregates containing both dyes generate a FRET signal. The resulting aggregates were then incubated with surface-immobilised dye-filled vesicles to quantify aggregate-vesicle colocalization. (**b, c**) Colocalization of vesicles and FRET-positive aggregates at 1 nM (B) and 200 pM (C) monomer-equivalent concentrations. Vesicles were excited at 488 nm and fluorescence was collected between 510-540 nm (green channel). FRET-positive aggregates were detected using 561 nm excitation and emission between 660-710 nm (magenta channel). (**d**) Distributions of $Ca^{2+}$ influx events measured at different aggregate concentrations (aggregation time point: 85 min, unlabelled). The distributions corresponding to Class A and Class B populations remain unchanged as aggregate concentration decreases. **(e)** Quantification of the total number of events leading to $Ca^{2+}$ influx at different aggregate concentrations, showing a linear dependence on aggregate concentration. **(f)** Total $Ca^{2+}$ influx summed across all vesicles at different aggregate concentrations, which also scales linearly with aggregate concentration.

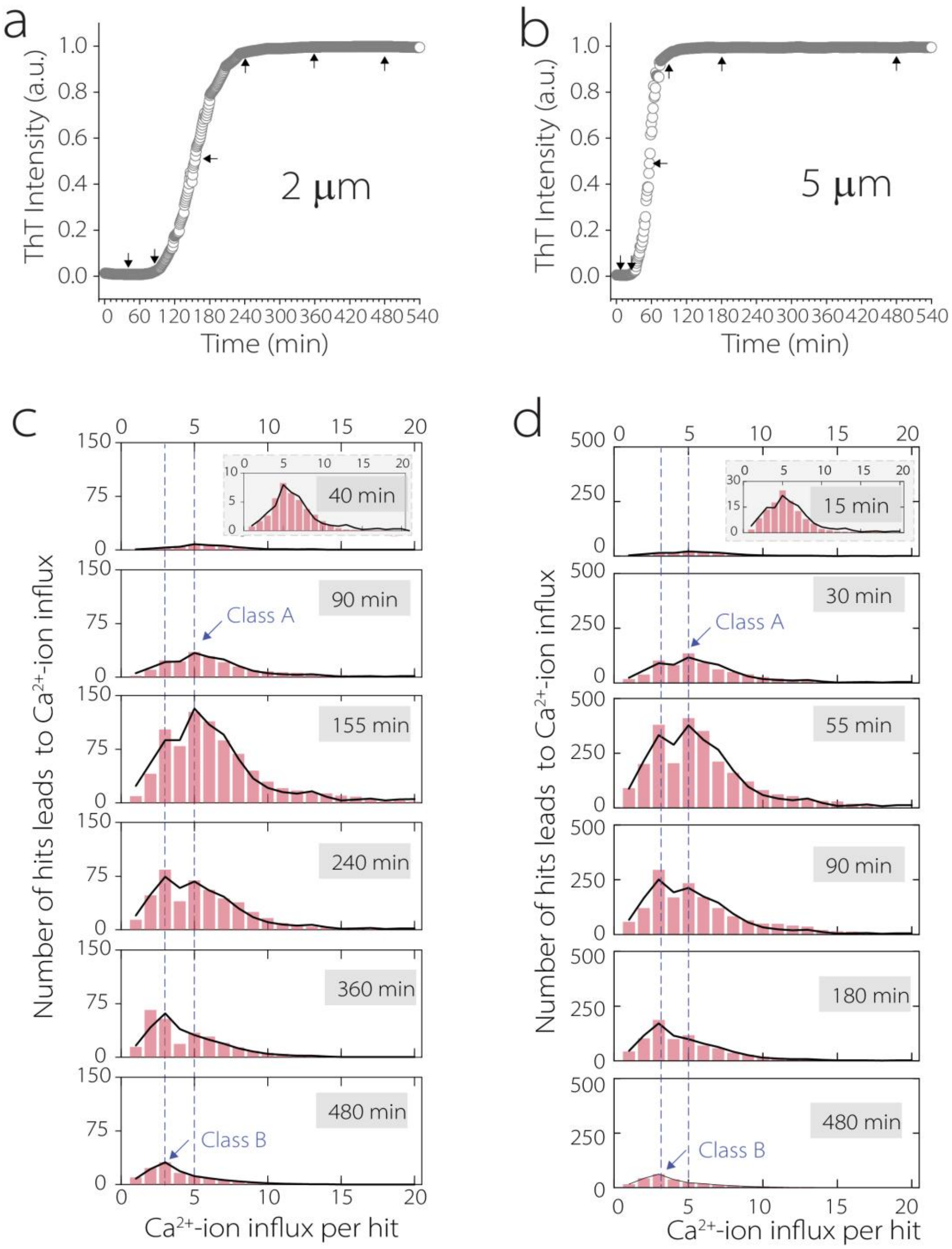


**Supplementary figure 4. (a, b)** ThT aggregation kinetics of Aβ42 at 2 μM (A) and 5 μM (B). Arrows indicate time points selected for downstream analysis. (**c, d)** Single-vesicle $Ca^{2+}$ influx distributions measured at corresponding time points for 2 μM (C) and 5 μM (D) Aβ42. Two dominant populations of intermediates were consistently observed for both concentrations.

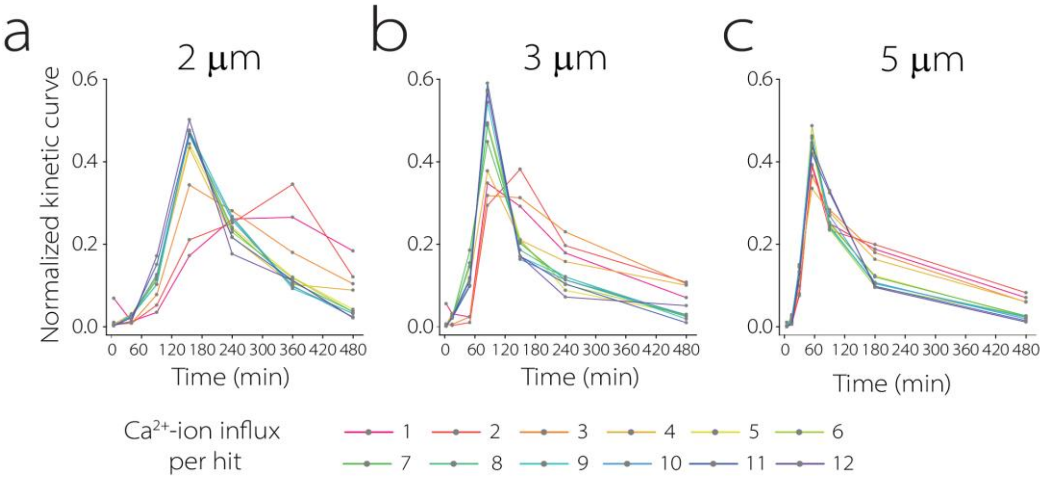


**Supplementary figure 5 (a-c).** Normalized kinetic curves illustrating the temporal evolution of Class A and Class B species at different Aβ42 concentrations **(a)** 2, **(b)** 3 and **(c)** 5 μm

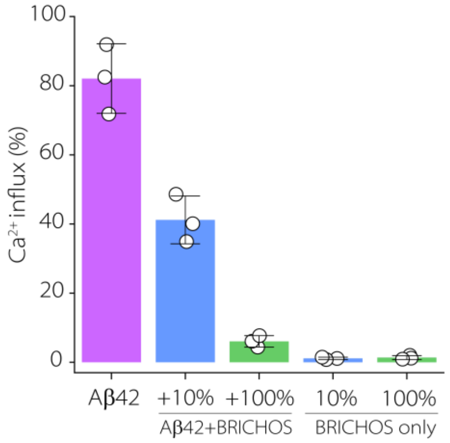


**Supplementary figure 6.** BRICHOS R221E does not disrupt vesicle membranes. Samples contained Aβ42 aggregates alone (85 min), Aβ42 aggregates in the presence of BRICHOS R221E (10% (105 min) and 100% (275 min) molar equivalents relative to Aβ42), or BRICHOS R221E alone (10% (105 min) and 100% (275 min) (Mean ± SD).

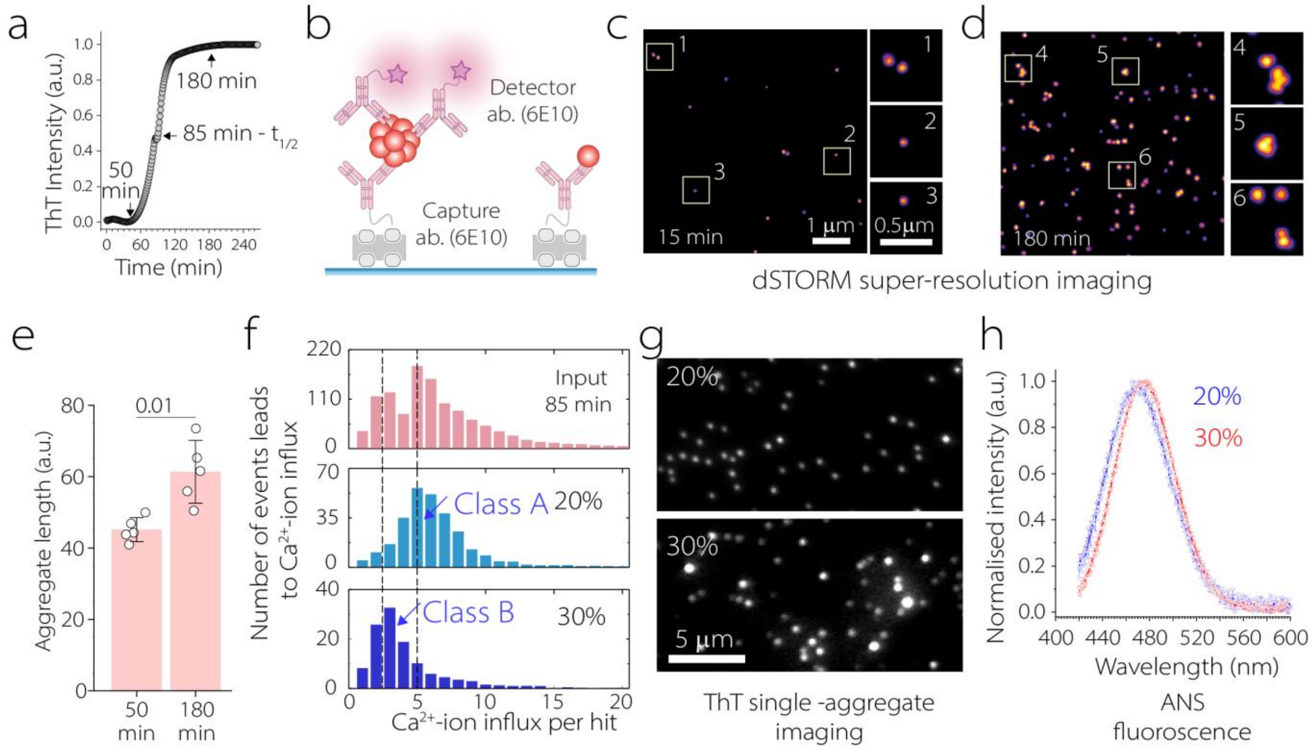


**Supplementary figure 7.** (**a**) ThT aggregation of 3 µM of Aβ42, (**b**) Schematic of the single-aggregate imaging assay. Aβ aggregates are captured on a surface using immobilised capture antibodies (6E10) and detected using Alexa Fluor 647 labelled detector antibodies (6E10), enabling imaging of individual Aβ assemblies, but not unaggregated monomers. (**c-d**) Representative dSTORM images of Aβ assemblies collected at 15 min and 180 min during aggregation. Insets show magnified examples of individual aggregates. (**e**) Quantification of aggregate size distribution measured from single-aggregate imaging. (**f**) $Ca^{2+}$ influx distributions measured for unfractionated aggregates collected at half time of the aggregation (top) and for aggregates isolated from 10-20% and 30-40% sucrose fractions. (**g**) Representative images of single aggregates present in the 10-20% and 30-40% sucrose fractions. **(h)** Normalized ANS fluorescence spectra of the 10-20% and 30-40% sucrose-gradient fractions, showing differences in surface hydrophobicity between the enriched aggregate populations.

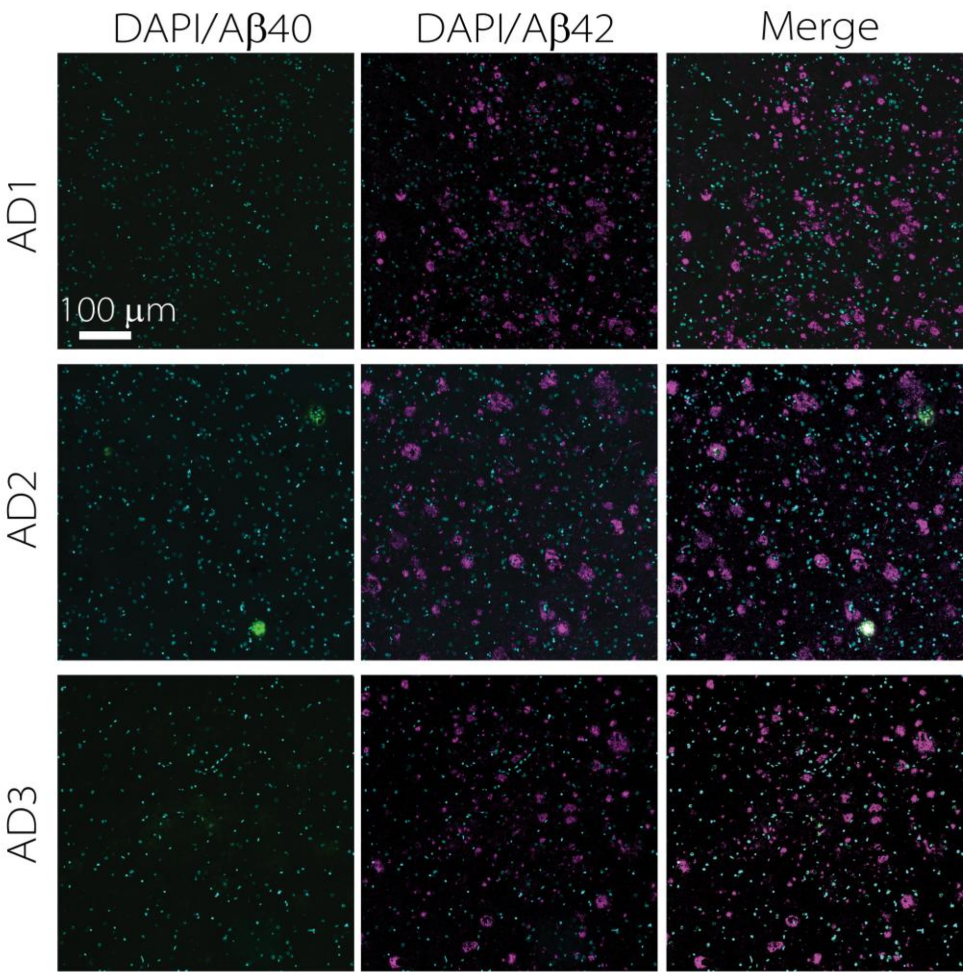


**Supplementary figure 8.** Immunohistochemistry of frontal cortex from three AD cases stained with the Aβ40-specific antibody (EPR23712-2) (green), Aβ42-specific antibody 21F12 (purple), and DAPI (cyan).

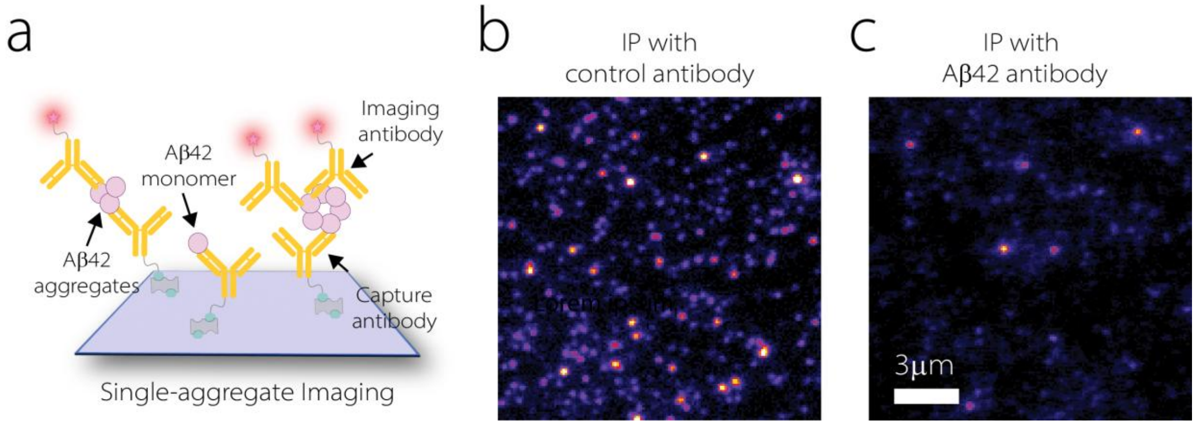


**Supplementary figure 9. (a)** Schematic of single-aggregate imaging in which both capture and detection are performed with the monoclonal antibody 21F12, allowing selective visualization of Aβ42 aggregates while excluding monomers. **(b, c)** Single-molecule fluorescence images of immunoprecipitation (IP) from AD brain extract using an isotype control antibody (B) or the Aβ42-specific antibody 21F12(C).

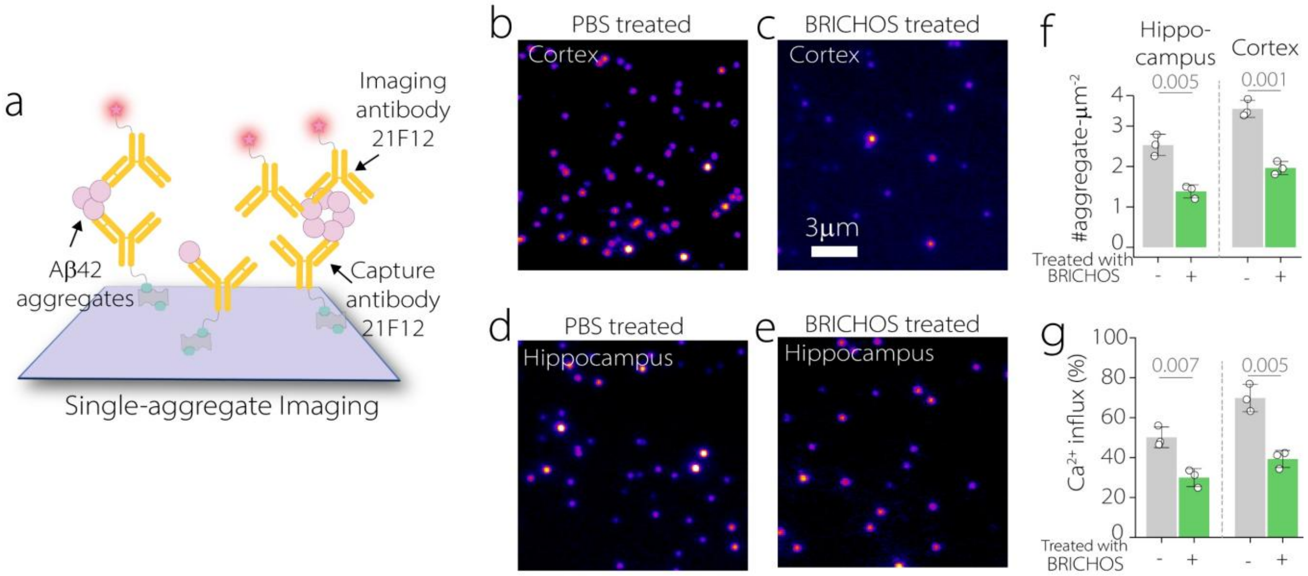


**Supplementary figure 10. (a)** Schematic of single-aggregate imaging using the monoclonal antibody 21F12. **(b-e)** Representative fluorescence images of immunoprecipitation (IP) from cortex (b, c) and hippocampus (d, e) of PBS-treated (b, d) and BRICHOS-treated (c, e) mice. **(f-g)** The number of aggregates (F) and their membrane permeabilization ability (G) measured by $Ca^{2+}$ influx was markedly reduced in hippocampal and cortex samples following BRICHOS treatment.

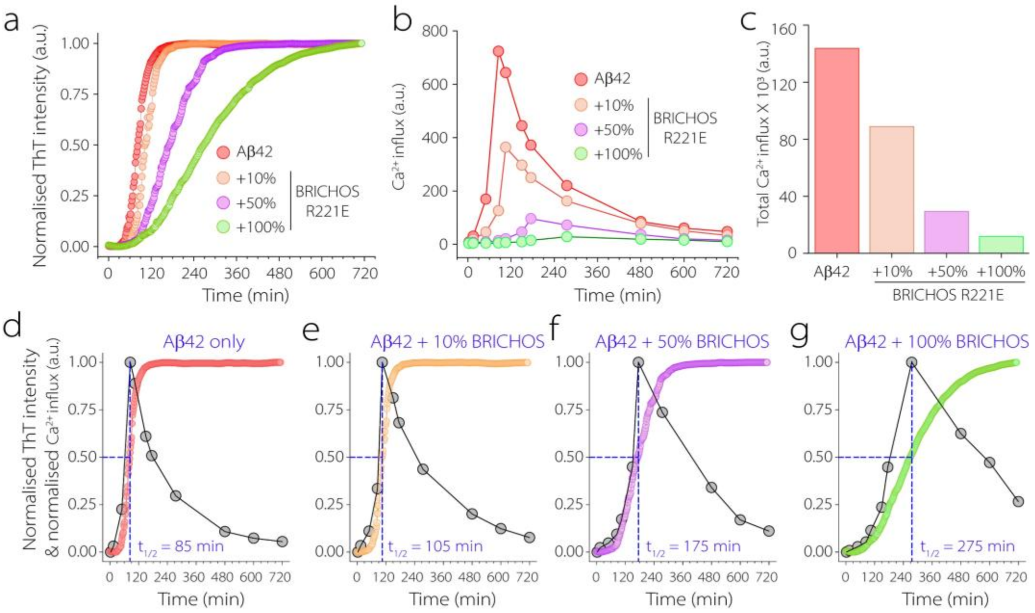


**Supplementary figure 11. (a)** Normalized ThT aggregation kinetics of Aβ42 alone or in the presence of increasing BRICHOS R221E concentrations. **(b)** $Ca^{2+}$ influx measured over the same aggregation time course using the vesicle assay. **(c)** Total $Ca^{2+}$ influx showing a dose-dependent reduction with increasing BRICHOS conc. **(d-g)** Overlay of normalized ThT aggregation kinetics and membrane-damaging activity for Aβ42 with 0% (D), 10% (E), 50% (F), or 100% (G) BRICHOS R221E. Dashed lines indicate the ThT half-time for each condition. Circles show experimental data points and lines are included as visual guides.

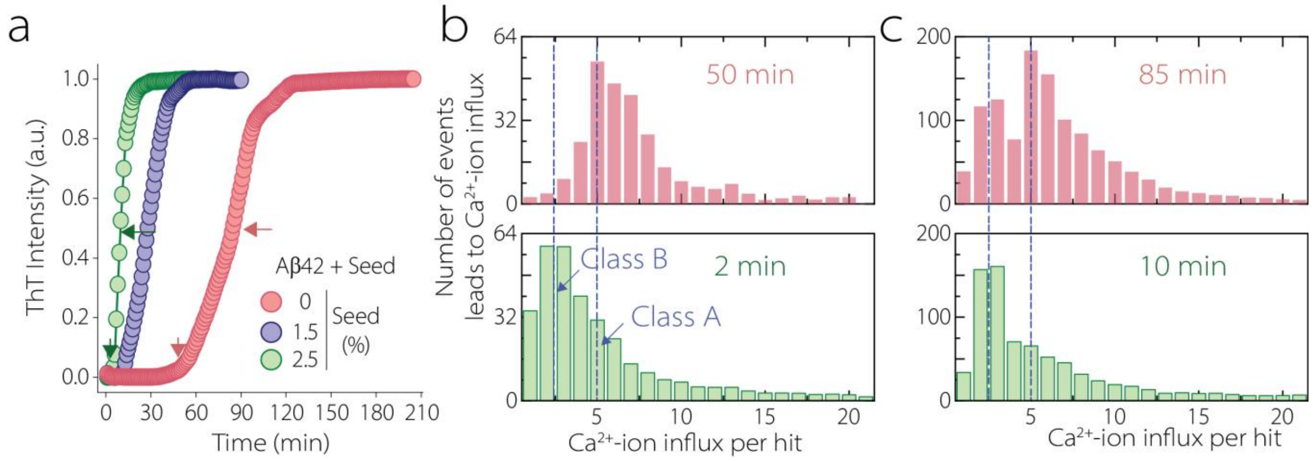


**Supplementary figure 12. (a)** ThT aggregation kinetics of Aβ42 in the absence or presence of 1.5% or 2.5% pre-formed fibril seeds. Arrows indicate the time points analysed. **(b-c)** $Ca^{2+}$ influx per hit distributions for unseeded and 2.5% seeded Aβ42 aggregation at the indicated time points.

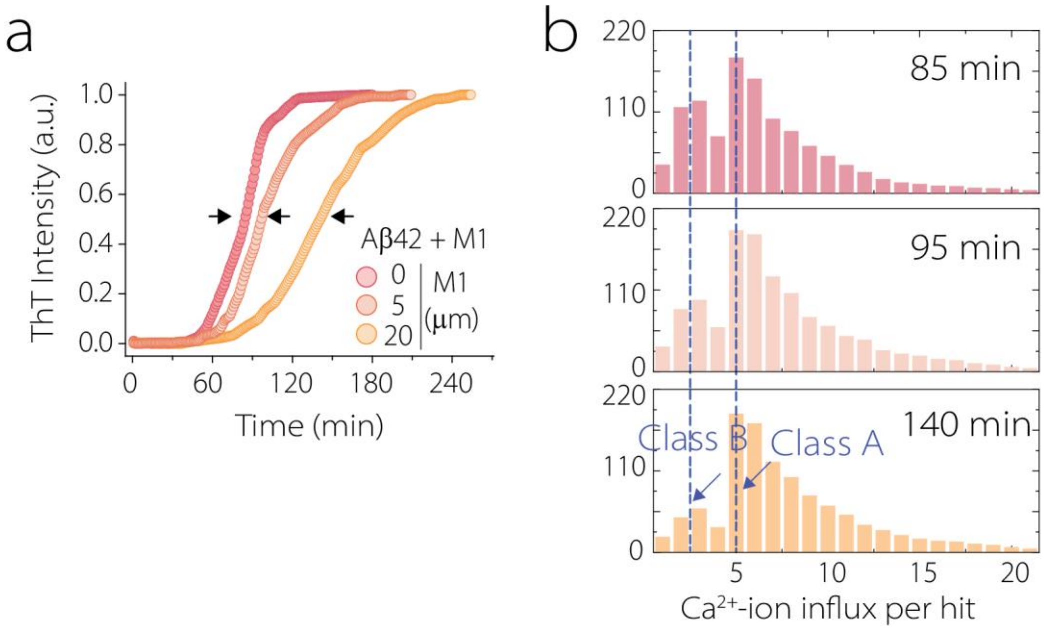


**Supplementary figure 13. (a)** ThT aggregation of Aβ42 in the absence or presence of increasing concentrations of M1. Arrows indicate the aggregation midpoints ($t_{1/2}$) analysed. **(b)** $Ca^{2+}$ influx per hit distributions measured at the corresponding midpoints for each aggregation.

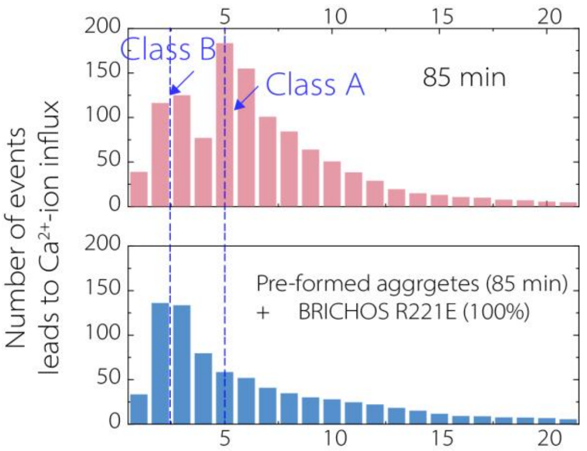


**Supplementary figure 14.** $Ca^{2+}$ influx per hit distributions for Aβ42 collected at the aggregation midpoint (85 min), before and after incubation with 100% molar equivalent of BRICHOS R221E monomer.

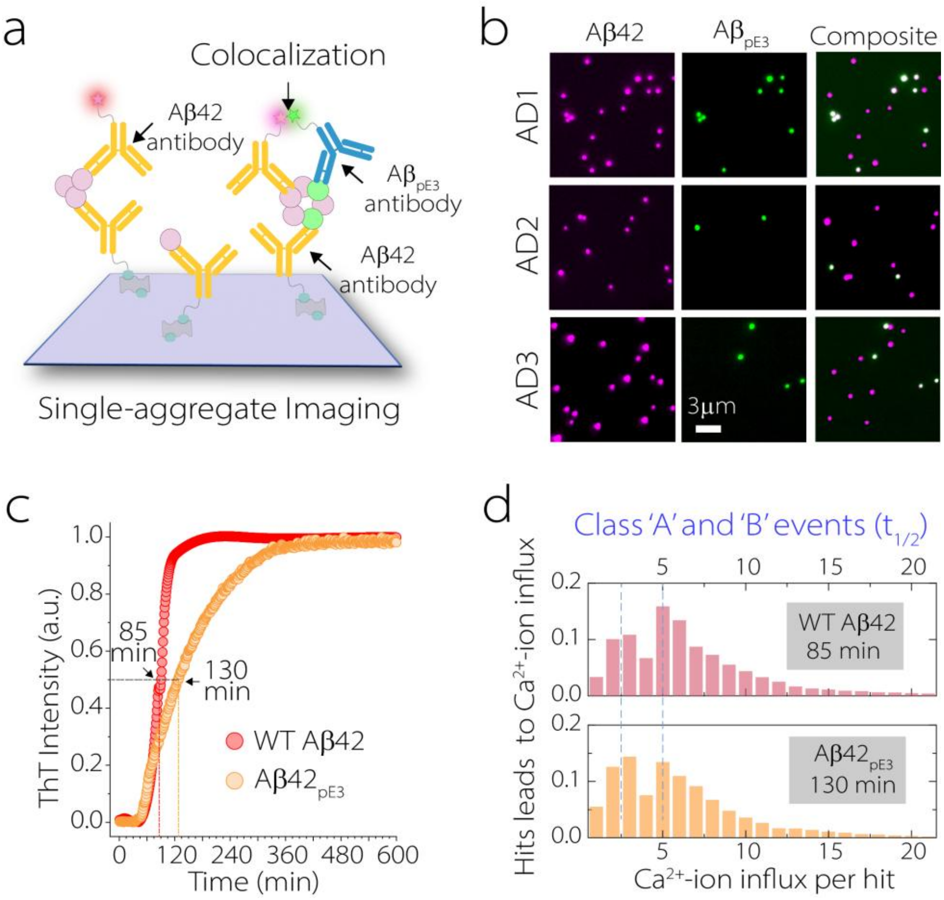


**Supplementary figure 15. (a)** Schematic of single-aggregate imaging, where Aβ42 aggregates are captured with 21F12 and imaged with antibodies against Aβ42 and $Aβ_{pE3}$. **(b)** Representative single-molecule fluorescence images from AD brain extracts showing aggregates positive for Aβ42 (magenta), $Aβ_{pE3}$ (green), and colocalization (white). **(c)** ThT aggregation kinetics of WT Aβ42 (red) and $Aβ_{pE3}$ (orange), showing a prolonged lag phase for $Aβ_{pE3}$. **(Dd)** Single-vesicle $Ca^{2+}$ influx distributions at the aggregation midpoint ($t_{1/2}$) reveal two populations of intermediates for both WT Aβ42 and $Aβ_{pE3}$, with comparable membrane-permeabilizing potency.

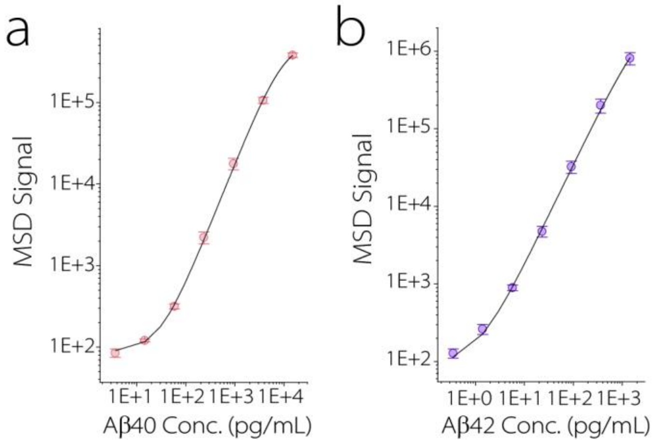


c

| | Aβ40 | Aβ42 |
|---|---|---|
| AD1 | 3.8 ± 0.6 | 55.4 ± 5.8 |
| AD2 | 5.7 ± 0.9 | 70..9 ± 6.9 |
| AD3 | 4.3 ± 0.3 | 58.7 ± 4.3 |

**Supplementary figure 16. (a, b)** Calibration curves for Aβ40 (A) and Aβ42 (B) generated using the Meso Scale Discovery (MSD) platform. **(c)** Measured concentrations of Aβ40 and Aβ42 in brain homogenates from three AD cases (ng/mL).

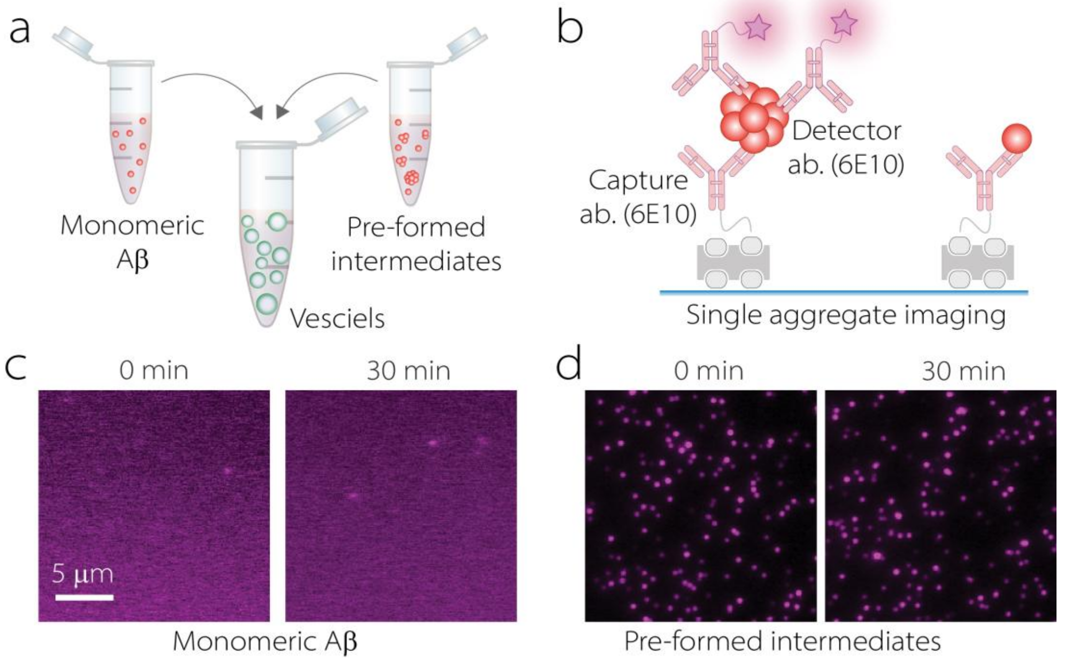


**Supplementary figure 17. (a)** Monomeric Aβ and pre-formed Aβ aggregates were incubated with dye-filled vesicles for 30 min at 37 ºC separately and then analysed using a single-aggregate imaging assay, in which aggregates are captured and detected using 6E10 antibody. (**b**) Representative single-aggregate images (contrast enhanced) of monomeric Aβ before (0 min) and after incubation with vesicles (30 min). No detectable aggregates are observed under either condition. (**c**) Representative single-aggregate images of pre-formed Aβ aggregates before (0 min – contrast enhanced) and after incubation with vesicles (30 min). The number of detected aggregates remains comparable following vesicle incubation.

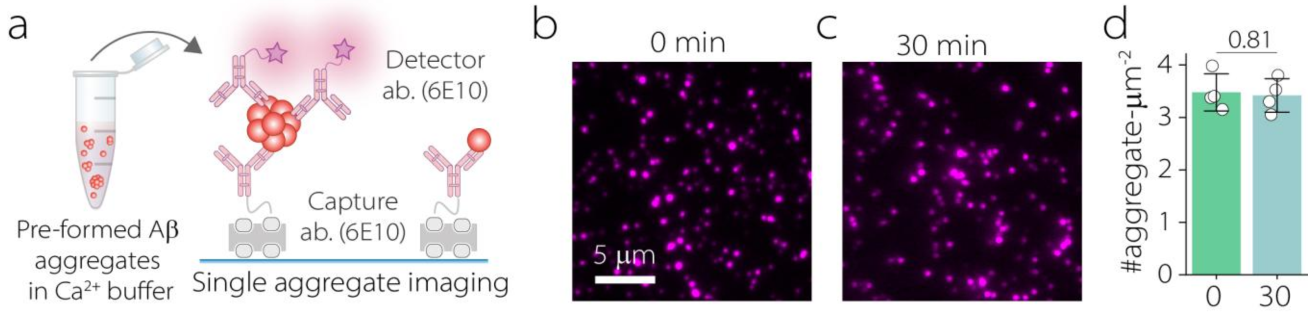


**Supplementary figure 18. (a)** Pre-formed Aβ aggregates were incubated in $Ca^{2+}$-containing buffer and analysed using a single-aggregate imaging assay. (**b**) Representative single-aggregate images of Aβ before and after incubation in $Ca^{2+}$-containing buffer. (**c**) Quantification of total Aβ aggregate counts measured by single-aggregate imaging before and after incubation in $Ca^{2+}$ containing buffer, showing no measurable change in aggregate abundance.

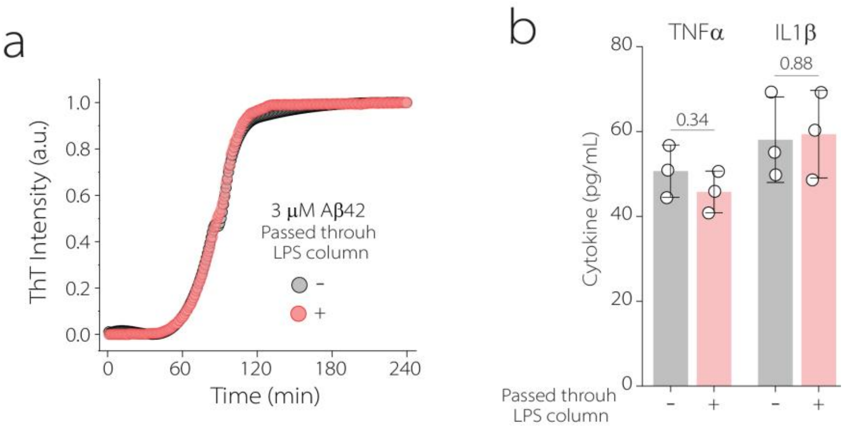


**Supplementary figure 19. (a)** ThT aggregation kinetics of recombinant Aβ42 before (grey) and after (red) endotoxin removal using an endotoxin removal spin column. **(b)** Astrocyte cytokine secretion following treatment with Aβ42 aggregates generated before (grey) or after (red) endotoxin removal. Cytokine secretions was similar under both conditions, indicating that the observed effects are not due to endotoxin contamination (Mean ± SD).

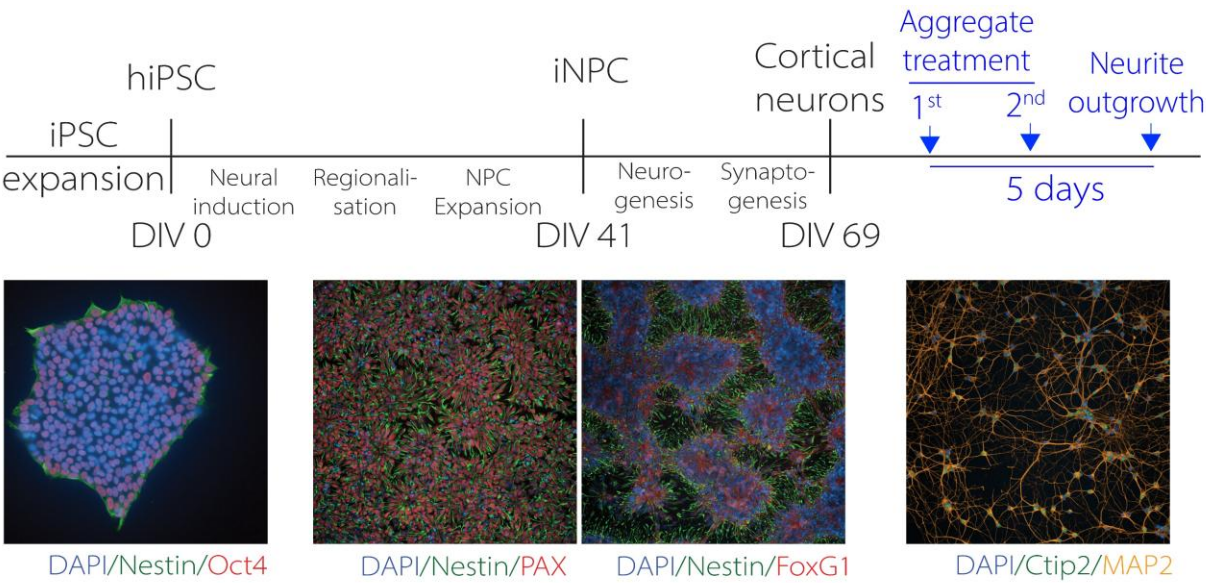


**Supplementary figure 20.** Differentiation timeline for generating cortical neurons from iPSCs via neuronal progenitor cells (iNPCs). Representative immunocytochemistry images illustrate the sequential stages of differentiation, showing iPSCs, iNPCs, and post-mitotic cortical neurons.

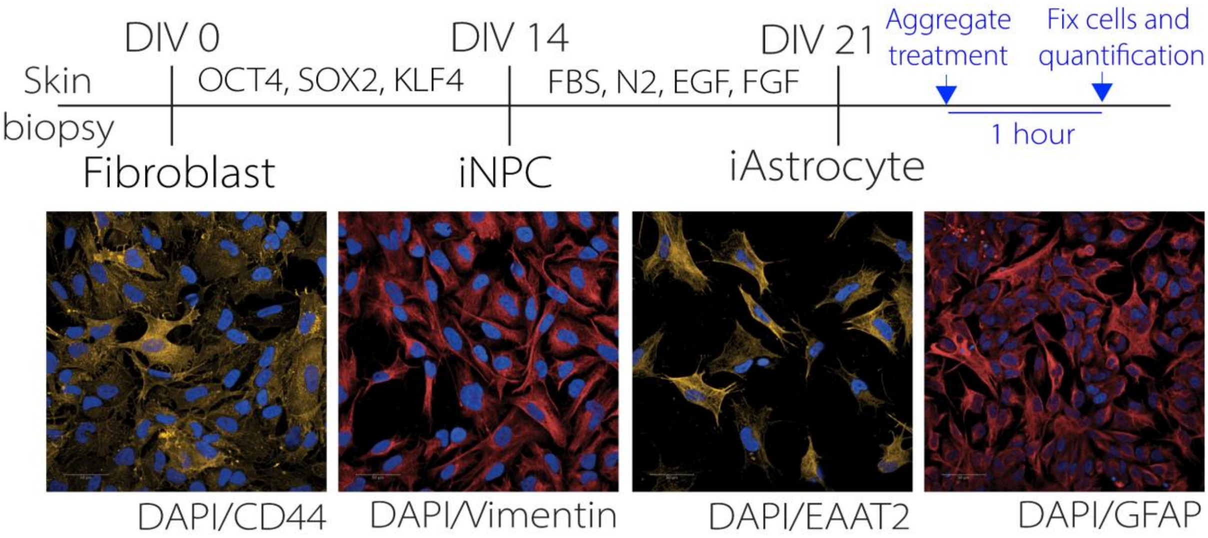


**Figure S21.** Differentiation protocol to generate iAstrocytes. Representative immunocytochemistry images showing expression of astrocyte markers CD44, Vimentin, EAAT2, and GFAP.
